# Special Nuclear Layer Contacts Among Starburst Amacrine Cells in the Mouse Retina

**DOI:** 10.1101/2022.08.28.504668

**Authors:** Shang Mu, Nicholas L. Turner, William M. Silversmith, Chris S. Jordan, Nico Kemnitz, Marissa Sorek, Celia David, Devon L. Jones, Doug Bland, Merlin Moore, Amy Robinson Sterling, H. Sebastian Seung, the Eyewirers

## Abstract

We observed novel classes of cell-cell contacts between retinal starburst amacrine neurons, from finely detailed morphological reconstructions of cells from an electron microscopic image volume of a mouse retina. These contacts have peculiar morphological patterns and traits, different among the respective On and Off starburst amacrine subpopulations, but both occur within the soma layers as opposed to their regular laminae of contact within the inner plexiform layer.

## INTRODUCTION

An abundant and well studied cell type in the mammalian retina is the starburst amacrine cells (SACs). They are so named by their characteristic starburst shape of their dendritic trees (Famiglietti, 1983; Tauchi and Masland, 1984). They come in two homologous subgroups: one group have their cell bodies in the ganglion cell layer of the retina and primarily respond to bright light stimuli and are called On SACs, while the other group have cell bodies in the inner nuclear layer and are better associated with stimuli’s transition from bright to darkness, hence named Off SACs. Initially identified as the acetylcholine synthesizing cells in the retina (Masland and Mills, 1979; Masland et al., 1984; Tauchi and Masland, 1984), these cells have well known identifying molecular labels (Wei and Feller, 2011; Ray et al., 2018), and their dendrites are tightly stratified such that SACs are the single most used location-referencing landmarks for studying cells in the context of the inner plexiform layer (IPL) of the retina (e.g., Sanes and Masland, 2015; Grünert and Martin, 2020).

In densely reconstructing and examining 199 starburst cells from an electron-microscopically (EM) imaged volume of a mouse retina patch (Briggman et al., 2011) in similar manners as we have reported before (Kim et al., 2014; Greene et al., 2016; Bae et al., 2018), we discovered special contact patterns.

Particularly, these contacts are on the cell bodies of other starburst cells (of the same On/Off subgroup). This is surprising because SACs have their dendrites tightly stratified in the IPL, and normally make synapses and contacts onto each other and with other types of cells via their dendrites within the IPL, as opposed to in the nuclear layers. We found peculiar morphological characteristics specific to each of the two respective subpopulations.

## RESULTS

### Ascending climbing dendrites of Off SACs

We discovered a novel class of contacts, each between a dendritic termination of one Off SAC and the soma and/or proximal dendrites of another Off SAC. These terminating dendrites veer towards the inner nuclear layer (INL), in many cases traveling almost parallel to the light axis (Fig 1A). This is unexpected because Off SAC dendrites normally stratify at a particular depth in the inner plexiform layer (IPL) and also terminate at that depth. The terminating dendrite often travels in contact with a proximal dendrite of the partner Off SAC, and if it reaches the partner cell’s soma, it typically spreads to a lump as it terminates (Fig 1B,C,D; Supplementary Fig 2). Most Off SAC cells (59 out of 96) in our dataset display this type of outbound and/or inbound contacts with one or more other Off SAC cells. Within these 59 cells, 38 radiate as many as 3 contacts each (1.3 ± 0.6, mean ± s.d.) to others, and 39 somas receive as many as 4 (1.3 ± 0.6, mean ± s.d.) contact patches from other cells.

**Figure 1:**
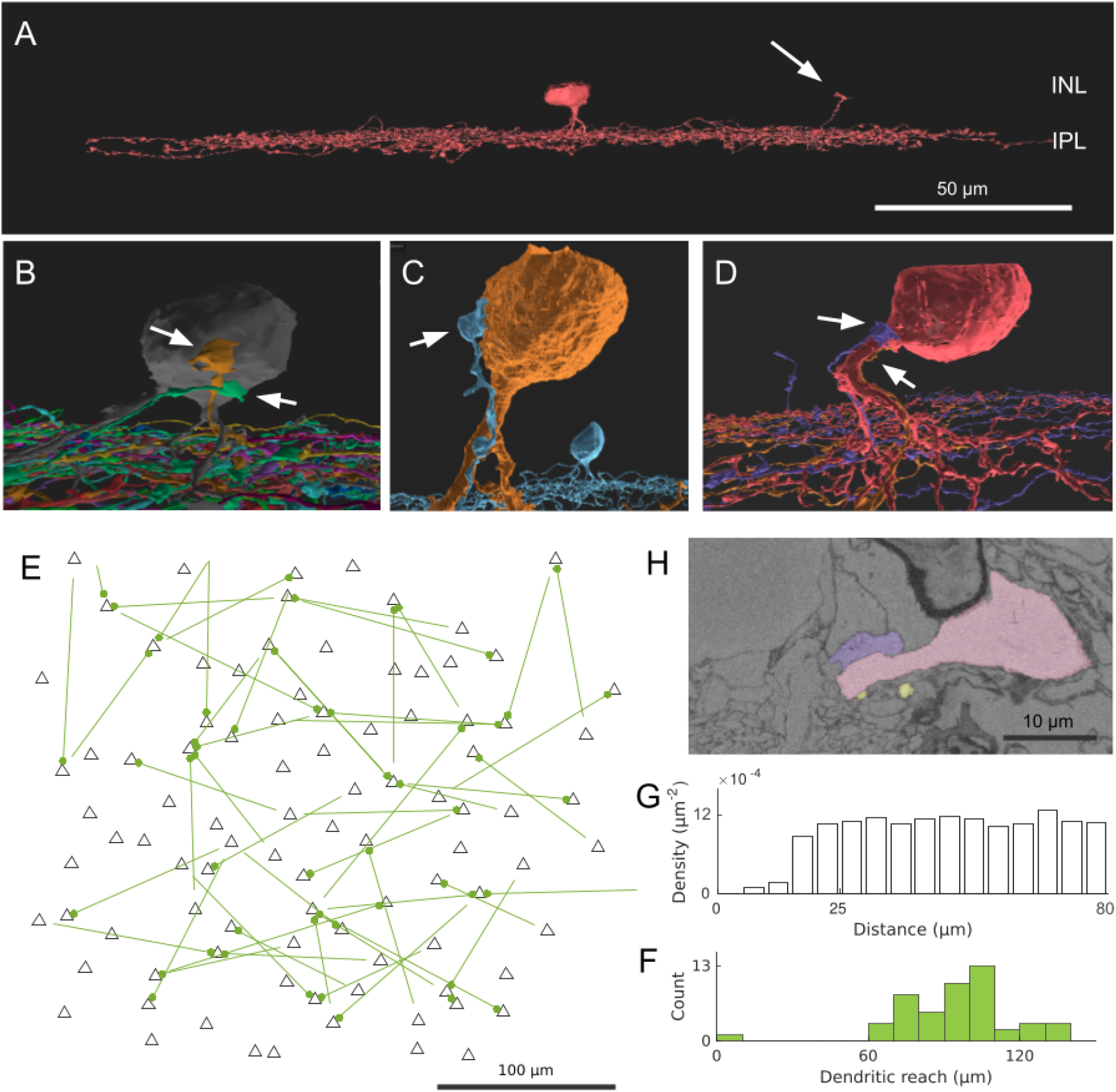
Off SAC contact pattern. **(A)** Tangential view of a 3D reconstructed Off SAC. An ascending dendrite (arrow) veers off from the dendritic stratification and into the inner nuclear layer (INL). **(B**,**C**,**D)** In each case of these 3D cell reconstructions, attached onto the soma or basal dendrite of an Off SAC we see ascending dendrites (arrows) from other Off SACs like in (A), usually climbing along the perisomatic dendrite. **(E)** Spatial distribution of the somatic origin and attachment points of these ascending dendrites in the retina patch (flat-mount view). Each line represents the dendritic branch starting from the originating cell’s soma (bare end of the line) and grasping onto the targeted cell’s perisomatic membrane (bulged end of the line as a dot); black triangles are soma locations of all reconstructed Off SACs with soma inside the retina patch. **(F)** Histogram showing the distribution of these ascending dendrites’ dendritic reach, defined as the planar distance from the ascending dendrite’s originating soma to the soma where it terminates. Minimum, quartiles, and maximum: 9, 81, 97, 104, 137 μm. **(G)** The density recovery profile, for all Off SAC somas regardless of any ascending dendrite contact or not, defined as the average density of somas at given distances from any given soma. **(H)** A mini region of the plasma-membrane-stained retina sample, shown as a sectional electron micrograph near the locations pointed to in (D), overlaid with the respective reconstructed cells’ colors matching panel (D). **Scale bars:** 50μm (A); 100μm (E); 10μm (H).

While no obvious pattern was seen (Fig 1E) in these perisomatically contacting Off-SAC to Off-SAC branches, we do see that the two cells in each contacting pair are rarely close to each other in terms of their somatic locations (Fig 1F), facilitated by the fact that in the flat-mount planar view these contacts are often near or at the most distal end of dendrites. In the 52 pairs of contacts we observed, only in one single case was the dendrite reaching for the SAC soma nearest to the originating soma, all other pairs were more than 60μm apart (Fig 1F). In comparison, the distribution of all Off SAC somas represented in the form of the density recovery profile (Rodieck, 1991), had already reached plateau at a distance of about 20-25μm (Fig 1G), indicating the closest SAC neighbor of a SAC was almost always less than 20μm away.

### Direct contacts and short processes bridging On SAC somata

We also discovered a novel class of contacts between somas of On SACs. In a prior study of SAC populations, Whitney et al. (2008) reported higher number of “close-neighbor pairs” in the ganglion cell layer (On SACs) than in the inner nuclear layer (Off SACs). Consistent with this observation, we found On SACs often pairing up next to each other in the ganglion cell layer (Fig 2A,B). Unexpectedly we noticed the pairs form intertwined short twigs at the contact in between the two somas (Fig 2B). In our specimen, 37 out of the 103 On SACs formed 19 adjoining pairs, with 14 pairs judged to have directly abutting somas (Fig 2A), and the remaining 5 pairs having dedicated short branch(es) reaching between the two somas from within the ganglion cell layer (Fig 2C). We consider two cells as a pair only if there exists direct contact of one of these two preceding forms within the ganglion cell layer between the two 3D-reconstructed cells. All pairs have flat-mount center-to-center soma distances within 17μm, and those for directly abutting pairs are all within 13μm (9 ± 2, mean ± s.d.). In comparison, the soma diameter of SACs measured under light microscopy are reported to be 10 (Farajian et al., 2004; Whitney et al., 2008) to 11 μm (Kay et al., 2012). All pairs exhibit twigs intertwined to various degrees, with the least prominent form being short stubs protruding from the cell body, hugging or protruding into the other cell body (Supplementary Fig. 1).

Whitney et al. (2008) argued that “close-neighbor pairs” were formed by cells displaced during development by fascicles of optic axons and retinal vasculature, from their original mosaic-proper locations. However, we have seen an example pair of two On SAC somata straddle an optic nerve fascicle, and also still have a dedicated short process bridging them (Fig 2C). Optic nerve fascicles can be a hindrance of forming pairs of cells abutting each other, and is therefore unlikely the cause of such formations. Another pair wrapped around a blood vessel, covering about 200 degrees of the blood vessel’s cross-sectional circumference. The blood vessel failed to cleanly separate the two somas which remain “touching hands” (data not shown). Combined with the intertwined twig-like structures present in all pairs, this suggests the close proximity and physical contact may carry concrete functional or developmental purpose. Occurrence of pairs was also reported in the rabbit in the ganglion cell layer (On SACs) and not in the inner nuclear layer (Off SACs) (Brandon, 1987), and similar higher rate of occurrence in the ganglion cell layer can be observed from published figures and images for cat retina (Fig. 4 in Vaney, 1990) and rabbit retina (Vaney et al., 1981; Fig. 7 in Clements et al., 2017).

**Figure 2:**
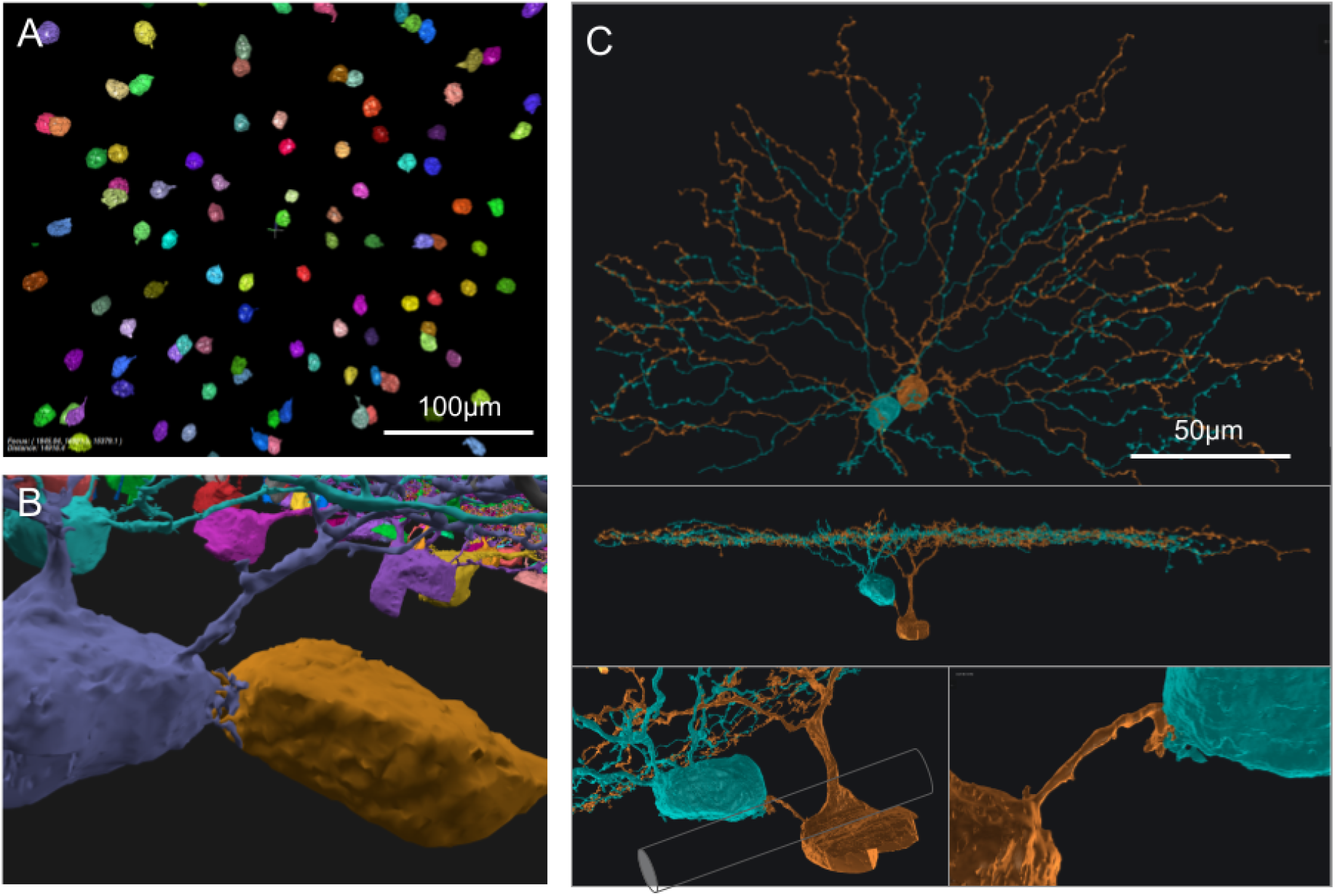
On SAC distribution and contacts. **(A)** Distribution of On SAC somas (colored objects are the 3D reconstructed full or partial somas) in a flat-mount view of the retina patch. **(B)** A pair of contacting On SACs form twigs at their soma contact (also in the background are somas of other On SACs, often incompletely reconstructed due to dataset boundaries). **(C)** A pair of On SACs that were next to each other in flat-mount view (top) did not have direct soma contact but were instead bridged by a short process between the somas (additional views in the middle and bottom panels). The two somas were separated by an axon bundle (cylindrical shape overlay within the bottom left panel) of retinal ganglion cells. **Scale bars:** 100μm (A); 50μm (C).

## DISCUSSION

### Off SAC perisomatic contacts

Ray et al. (2018) studied SAC development and reported that Off SACs establish dendrite-soma contacts during radial migration and assume transitory bi-laminar dendritic morphology consisting of a soma-layer lamina with soma-layer SAC-SAC contacts upon completion of migration. These soma-layer contacts, however, are mostly eliminated by day P3. It is possible the dendrite-soma contacts we observed are remnants of these developmental processes. The retina dataset we have is from a wild-type (C57BL/6) mouse of age P29 (Briggman et al., 2011). On the other hand, the perisomatic contacts we observed were almost never (1 out of 52, Fig 1F,E) between two close-by “neighboring” Off starburst cells. These ascending dendrites are usually far away from their originating somata, and are close to, or themselves are, the far terminating tip of the SAC dendrite carrying them. There are therefore usually several cell bodies of other starburst cells between the two cell bodies of the contacting pair of this kind. This is in contrast to the soma-layer contacts Ray et al. (2018) reported, which were exclusively between neighboring SACs.

Within individual starburst cells, different dendrites are known to function quite independently of each other in experiments examining light-evoked responses (Euler et al., 2002; Lee and Zhou, 2006; Morrie and Feller, 2018). The occurrence of our ascending-dendrite contacts becomes quite rare if we compare against the total number of distal dendrites rather than the number of cells, i.e., if we were to regard these dendrites just like other (independently functioning) distal dendrites and if they relay and compute just the same kind of information a regular SAC distal dendrite relays. This rareness and perceived “insignificant contribution” can call for an argument that light-invoked responses are less likely a functional target affected by these connections, except for perhaps long range interactions across the retina, or extremely local information where a single ascending dendrite should dominate all.

This known functional independence of individual dendrites pertains to light stimuli (with spatial details) but does not preclude potentially non-independent regulatory functions, for example, developmental purposes. The fact that these contacts are on the cell bodies hints at a more cell-centric function rather than a local dendritic-centric function.

Not all Off SAC cells were observed to have this class of contacts on them. However, with the extremely high coverage factor of starburst cells in the retina (exceeding 30, Keeley et al., 2007), just a small portion of these cells would already have the capacity to cover the entire retina.

### On SAC short bridging processes

In Ray et al. (2018), similar to Off SAC dendrite-soma contacts, On SACs also made soma layer projections between days P0 and P3 that contacted neighboring SAC somata. The transcellular signal Megf10 protein was found to promote the formation of dendritic sublayer within the inner plexiform layer and the elimination of arbor projections in soma layer by P3. The same protein additionally controlled development of proper mosaic spacing of somas beyond P3 (Kay et al., 2012; Ray et al., 2018). It is possible that the bridging processes we found were to help “push apart” neighboring somas before they eventually degenerate or retract, but such possibility is remote given that somata in these pairs we saw were all immediately adjacent to each other in the flat-mount projection view, being gross violations of the frequently acknowledged near-soma dead zone (Rodieck, 1991; Galli-Resta et al., 2000) or mosaic rule (Wässle and Riemann, 1978; Vaney et al., 1981). Abutting On SAC soma pairs were not specifically reported by Kay et al. (2012) but were indeed visible and frequent in its Fig. S3 for wild-type mice, making our observations consistent with theirs.

Ray et al. (2018) additionally observed single unbranched processes extending from the somata, ∼180° away from the IPL. These “180° arbors” were reported to be sometimes still present in P5 SACs, and were considered to be fundamentally different from the tangentially projecting soma-layer neurites in the developing retina. The bridging processes in our P29 retina are also unbranched and in principle could also potentially be related to this second class of unusual processes.

Our reconstruction is known to be incomplete within the nuclear layers due to the dataset boundary and because our automated convolutional neural network algorithm that facilitated our reconstruction were not especially well trained for the deep nuclear regions. The branches in both cases of the SAC subpopulations are thin and can be missed due to staining gaps or just proofreading oversight. For the On SACs it is possible and likely we missed some of the connecting branches especially overpassing the data boundaries at the ganglion cell layer. For the Off SACs it is also possible we missed certain contacting patches if these dendrites reach well into the inner nuclear layer. However, the novel contacts we found are still abundant and not unique cases of random mutations.

While it is not entirely surprising that electron microscopic reconstructions give finer views into the complex network of neuronal connections, we were nevertheless amazed by these novel contacts not reported by prior studies. We believe the reasons why these were never reported before are two fold: First, in light microscopy these cells need to be filled sparsely or differentially in order for these climbing dendrites and contacts to be seen, and staining efficiency and signal-to-noise ratio become limiting factors in recognizing these contacts at the far-tip. Electron microscopy does not have these limitations but were typically done only in tiny volumes. Second, neither light microscopy or electron microscopy was traditionally volumetric, and these contacts only became apparent when a fully 3D visualization is employed, advantageous compared to single section visualizations.

Due to historical reasons, this particular electron microscopy (EM) volume we used did not have intracellular staining and we were unable to identify these nuclear-layer contacts as synaptic or otherwise (Fig 1H). Further studies of the subcellular structure and molecular identities are warranted for these special SAC-SAC contact sites.

## ACKNOWLEDGEMENTS

We thank Kevin Briggman for providing the raw electron microscopic dataset, identified as e2198.

## SUPPLEMENTARY MATERIALS

### Supplementary Notes

This file includes a list of individual Eyewirers who contributed to the reconstruction of the starburst cells in this report.

### Supplementary Figures

This file includes additional examples and close-up views of the contact sites in both EM sections and 3D views.

## METHODS

### Neuron reconstruction

Neurons were reconstructed mostly via the online citizen science game Eyewire.org, as reported in previous publications (Kim et al., 2014; Greene et al., 2016; Bae et al., 2018), as well as part of newer campaigns in the game. Additional effort was also carried out to search for Off SAC characteristic patches and climbing dendrites on some somas where no incoming SAC contact had already been observed; when such patches or dendrites were found, they were inserted into the Eyewire system for reconstruction. A small number of these found instances were back traced to existing SAC reconstructions where the branches were previously missed or mistaken as reconstruction errors due to their unusual course of extension. A number of them resulted in reconstruction of full starburst cells that had not yet been reconstructed in the normal course of campaigns in the Eyewire game at the time.

## Density recovery profile of Off SAC somas (Fig. 1G)

We computed the density recovery profile (Rodieck, 1991) taking all Off SAC somas as reference points, inclusive of those closer to the dataset boundary.

We first computed all the pairwise flat-mount planar distance between Off SAC somas, and binned them into 5μm bins. Without normalization, this would give a traditional histogram plot. We then normalized each bin count by dividing it with the area of the rings at the given planar distance from SAC somas in the dataset (respecting dataset boundaries, see below), thereby obtaining the density recovery profile. Each pairwise distance is counted twice due to the symmetric relationship between the two members of each pair.

For inclusion of somas closer to the boundary, we did not use the method of correction factors (Rodieck, 1991) that uses “mean” effective sampling areas relying on an assumption of relatively uniform distribution of reference points, which would be entirely reasonable if the number of reference points was big enough. Instead, the concentric annuluses centered at each soma location were intersected with the bounding rectangle of the soma centers of these SACs, to produce the actual intersection areas which are used for the normalization described in the preceding paragraph.

## References

Bae, J. A., Mu, S., Kim, J. S., Turner, N. L., Tartavull, I., Kemnitz, N., et al. (2018). Digital Museum of Retinal Ganglion Cells with Dense Anatomy and Physiology. Cell 173, 1293–1306.e19. doi: 10.1016/j.cell.2018.04.040.

Brandon, C. (1987). Cholinergic neurons in the rabbit retina: dendritic branching and ultrastructural connectivity. Brain Res. 426, 119–130. doi: 10.1016/0006-8993(87)90431-8.

Briggman, K. L., Helmstaedter, M., and Denk, W. (2011). Wiring specificity in the direction-selectivity circuit of the retina. Nature 471, 183–188. doi: 10.1038/nature09818.

Clements, R., Turk, R., Campbell, K. P., and Wright, K. M. (2017). Dystroglycan Maintains Inner Limiting Membrane Integrity to Coordinate Retinal Development. J. Neurosci. 37, 8559–8574. doi: 10.1523/JNEUROSCI.0946-17.2017.

Euler, T., Detwiler, P. B., and Denk, W. (2002). Directionally selective calcium signals in dendrites of starburst amacrine cells. Nature 418, 845–852. doi: 10.1038/nature00931.

Famiglietti, E. V., Jr (1983). “Starburst”amacrine cells and cholinergic neurons: mirror-symmetric ON and OFF amacrine cells of rabbit retina. Brain Res. 261, 138–144. Available at: https://www.sciencedirect.com/science/article/pii/0006899383912933.

Farajian, R., Raven, M. A., Cusato, K., and Reese, B. E. (2004). Cellular positioning and dendritic field size of cholinergic amacrine cells are impervious to early ablation of neighboring cells in the mouse retina. Vis. Neurosci. 21, 13–22. doi: 10.1017/s0952523804041021.

Galli-Resta, L., Novelli, E., Volpini, M., and Strettoi, E. (2000). The spatial organization of cholinergic mosaics in the adult mouse retina. Eur. J. Neurosci. 12, 3819–3822. doi: 10.1046/j.1460-9568.2000.00280.x.

Greene, M. J., Kim, J. S., Seung, H. S., and EyeWirers (2016). Analogous Convergence of Sustained and Transient Inputs in Parallel On and Off Pathways for Retinal Motion Computation. Cell Rep. 14, 1892–1900. doi: 10.1016/j.celrep.2016.02.001.

Grünert, U., and Martin, P. R. (2020). Cell types and cell circuits in human and non-human primate retina. Prog. Retin. Eye Res., 100844. doi: 10.1016/j.preteyeres.2020.100844.

Kay, J. N., Chu, M. W., and Sanes, J. R. (2012). MEGF10 and MEGF11 mediate homotypic interactions required for mosaic spacing of retinal neurons. Nature 483, 465–469. doi: 10.1038/nature10877.

Keeley, P. W., Whitney, I. E., Raven, M. A., and Reese, B. E. (2007). Dendritic spread and functional coverage of starburst amacrine cells. J. Comp. Neurol. 505, 539–546. doi: 10.1002/cne.21518.

Kim, J. S., Greene, M. J., Zlateski, A., Lee, K., Richardson, M., Turaga, S. C., et al. (2014). Space–time wiring specificity supports direction selectivity in the retina. Nature 509, 331–336. doi: 10.1038/nature13240.

Lee, S., and Zhou, Z. J. (2006). The synaptic mechanism of direction selectivity in distal processes of starburst amacrine cells. Neuron 51, 787–799. doi: 10.1016/j.neuron.2006.08.007.

Masland, R. H., and Mills, J. W. (1979). Autoradiographic identification of acetylcholine in the rabbit retina. J. Cell Biol. 83, 159–178. doi: 10.1083/jcb.83.1.159.

Masland, R. H., Mills, J. W., and Hayden, S. A. (1984). Acetylcholine-synthesizing amacrine cells: identification and selective staining by using radioautography and fluorescent markers. Proc. R. Soc. Lond. B Biol. Sci. 223, 79–100. doi: 10.1098/rspb.1984.0084.

Morrie, R. D., and Feller, M. B. (2018). A Dense Starburst Plexus Is Critical for Generating Direction Selectivity. Curr. Biol. 28, 1204–1212.e5. doi: 10.1016/j.cub.2018.03.001.

Ray, T. A., Roy, S., Kozlowski, C., Wang, J., Cafaro, J., Hulbert, S. W., et al. (2018). Formation of retinal direction-selective circuitry initiated by starburst amacrine cell homotypic contact. Elife 7. doi: 10.7554/eLife.34241.

Rodieck, R. W. (1991). The density recovery profile: a method for the analysis of points in the plane applicable to retinal studies. Vis. Neurosci. 6, 95–111. doi: 10.1017/s095252380001049x.

Sanes, J. R., and Masland, R. H. (2015). The types of retinal ganglion cells: current status and implications for neuronal classification. Annu. Rev. Neurosci. 38, 221–246. doi: 10.1146/annurev-neuro-071714-034120.

Tauchi, M., and Masland, R. H. (1984). The shape and arrangement of the cholinergic neurons in the rabbit retina. Proc. R. Soc. Lond. B Biol. Sci. 223, 101–119. doi: 10.1098/rspb.1984.0085.

Vaney, D. I. (1990). Chapter 2 The mosaic of amacrine cells in the mammalian retina. Prog. Retin. Eye Res. 9, 49–100. doi: 10.1016/0278-4327(90)90004-2.

Vaney, D. I., Peichi, L., and Boycott, B. B. (1981). Matching populations of amacrine cells in the inner nuclear and ganglion cell layers of the rabbit retina. J. Comp. Neurol. 199, 373–391. doi: 10.1002/cne.901990305.

Wässle, H., and Riemann, H. J. (1978). The mosaic of nerve cells in the mammalian retina. Proc. R. Soc. Lond. B Biol. Sci. 200, 441–461. doi: 10.1098/rspb.1978.0026.

Wei, W., and Feller, M. B. (2011). Organization and development of direction-selective circuits in the retina. Trends Neurosci. 34, 638–645. doi: 10.1016/j.tins.2011.08.002.

Whitney, I. E., Keeley, P. W., Raven, M. A., and Reese, B. E. (2008). Spatial patterning of cholinergic amacrine cells in the mouse retina. J. Comp. Neurol. 508, 1–12. doi: 10.1002/cne.21630.

