## Supplementary Figures for "Special Nuclear Layer Contacts Among Starburst Amacrine Cells in the Mouse Retina"

**Supplementary Figure 1**
Additional examples of contacting On SAC somas

**Top row**: Electron microscopic sectional views of the contacting surface in different pairs. The contacting somas (asterisks) form protrusions and twigs, commonly at the the contacting surface boundary - intertwined (paired arrows) or hugging the partner soma (singular arrow), and sometimes in the middle of the contacting surface (arrowheads), with the last example being an "entangled ball" formed by the intertwined twigs right in the middle of a soma-soma contacting surface (paired arrowheads).

**Bottom row**: 3D perspective views showing the “hugging” type of protrusions at the boundaries of the main contacting surfaces.

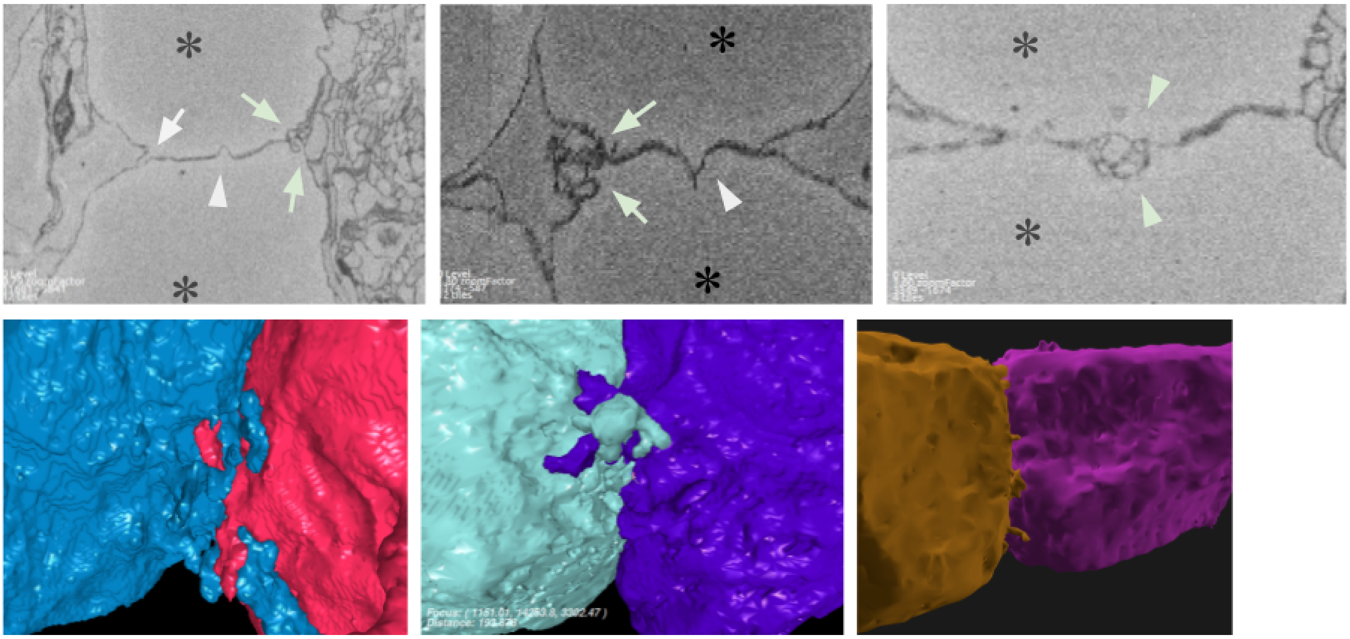

**Supplementary Figure 2**

Additional examples of ascending dendrite contacts on Off SAC somas (orthogonal 2D sectional views of the electron microscopic volume and color-matched 3D views from various directions)

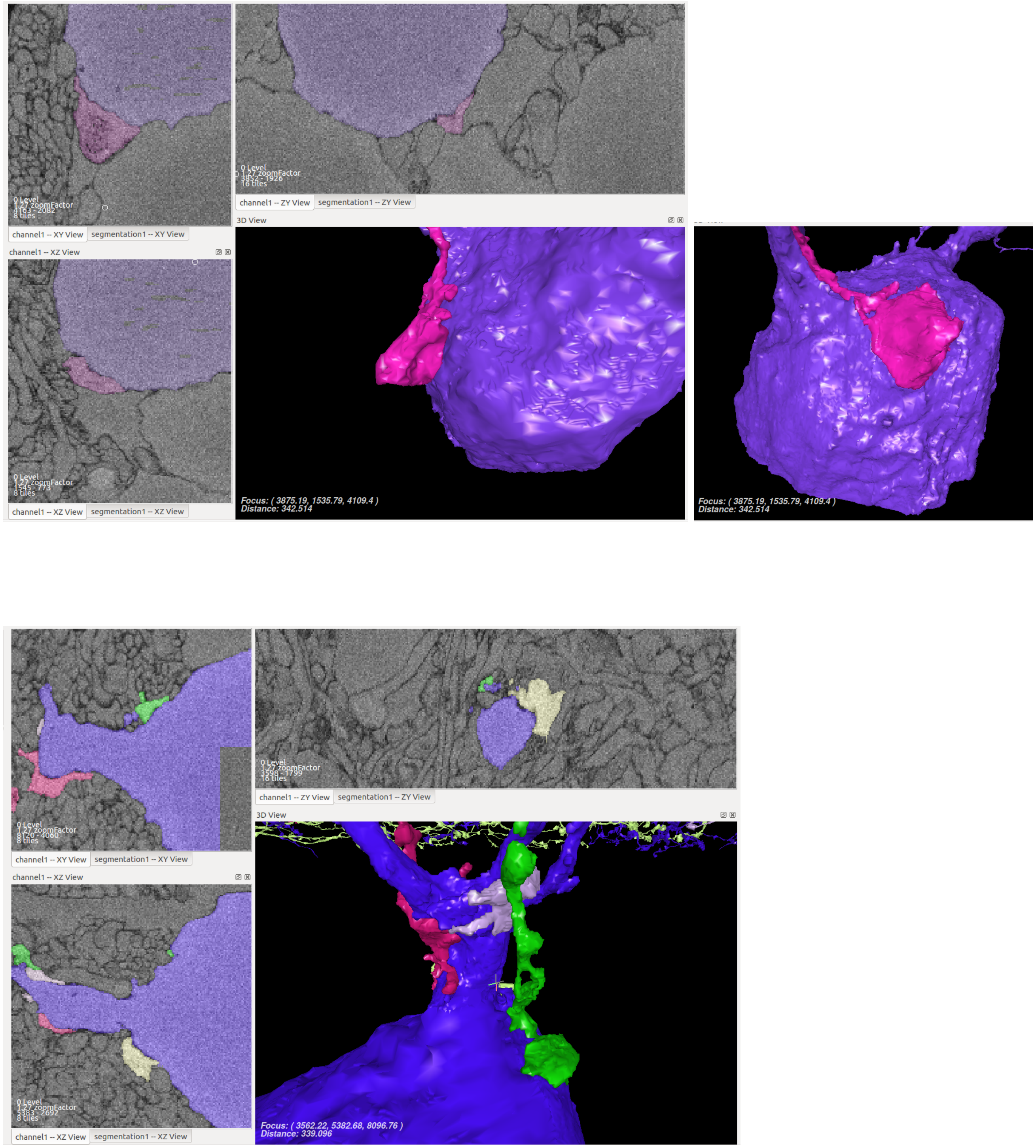

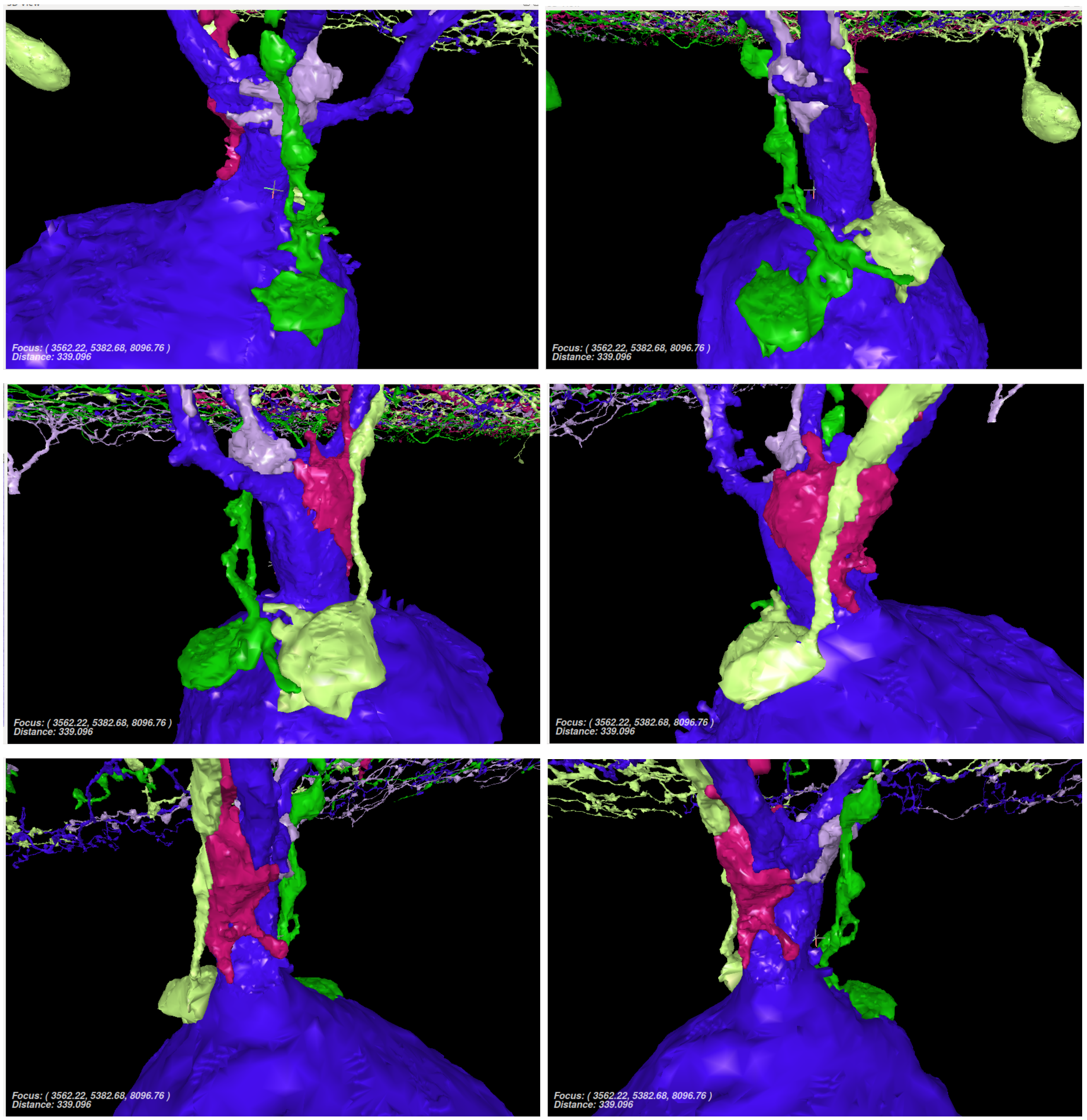
Focus: ( 3562.22, 5382.68, 8096.76 )
Distance: 339.096
Focus: ( 3562.22, 5382.68, 8096.76 )
Distance: 339.096
Focus: ( 3562.22, 5382.68, 8096.76 )
Distance: 339.096
Focus: ( 3562.22, 5382.68, 8096.76 )
Distance: 339.096
Focus: ( 3562.22, 5382.68, 8096.76 )
Distance: 339.096
Focus: ( 3562.22, 5382.68, 8096.76 )
Distance: 339.096
