## Supplementary Notes for "Special Nuclear Layer Contacts Among Starburst Amacrine Cells in the Mouse Retina"

The following Eyewirers took on leadership roles during the reconstruction of starburst amacrine cells:

- Mentors helped novice players to integrate into the community and improve their tracing skills: Nseraf, aesanta1, galarun, mamateresa, michellewooten, damocles357, sambob496, DannyS, jax123, IcyCuber, addieye, susi, LotteryDiscountz, kronnn, Emeraldstar, ronin, Varppi, b_hailey, jungmanp88, davidheiserca, superrogin, goodjelly, MaraTara, Andrearwen, tunerk7, reb1618, mam711, djajsl1234, scoobi, a5hm0r, Caffeine, lilmoo, Atani, dataminerstarr, LindseyAB, FishKAA, kinryuu, LCU, eldendaf, pkeoughan, skruffylooter, Queensalis, Manni_Mammut, jfine, nopasaran, dragonturtle, Byuda, Fabiett, wm05055, pokemontrainer, r3, Komorebi, nagilooh, Cliodhna, lgrochow, argento92, sjw3310, vitadolce5, randompersonjci, pollysingerman, gardenpea, aluorvats, falconheart, mkwak, m7md, antoineb, Omoshne, annkri, baraitalo, Frosty, Lochnivar, Kfay, JousterL, cerebralcortex, Peridot, Here_n_Now, Xnopyt, Baraka, juneyang2005, ggreminder, hewhoamareismyself Oppen_heimer, hawaiisunfun, NoelleO, LeeMo127, canigotothemoonpls GFani, cognaso, TheAtom417.
- Moderators helped mobilize the community and intervened in case of conflict between players: rinda, jinbean, galarun, susi, a5hm0r, sarabc, tek50, Nseraf, kronnn, DannyS, marhav, addieye, lilmoo, michellewooten, mamateresa, ronin, b_hailey, jax123, jungmanp88, LD2, LotteryDiscountz, angler.korea, aesanta1, pkeoughan, minsuyang, MaraTara, northingood, KateLee, darwinkim, ttore63, nspainter77, benficus, tunerk7, superrogin, xodhks9205, martyy_plp, djajsl1234, scoobi, goodjelly, Andrearwen, m7md, mam711, ehmilkyway, skruffylooter, LindseyAB, Sunnyway, Atani, reb1618, graigrai, kinryuu, lgrochow, Queensalis, r3, gracemansfield, Fabiett, lusiyan, nopasaran, tfrank, wm05055, Manni_Mammut, Cliodhna, vitadolce5, sjw3310, argento92, zRockeTz, Emeraldstar, Lunar_Dust, aluorvats, baraitalo, lllhv66, hanm0, mans0930, Omoshne, annkri, antoineb, dragonturtle, Komorebi, KrzysztofKruk, JousterL, Zeekza, Xnopyt, hawaiisunfun, NoelleO, Caffeine, LynneC, falconheart, Kfay, hewhoamareismyself Oppen_heimer, nagilooh, damocles357, THORR, GFani, SpookyGrowly.
- Scouts reported possible reconstruction errors: tschwach, faunhaert, aluorvats, Fritzchen, Cliodhna, hawaiisunfun, Fess, djajsl1234, wm05055, Omoshne, mcsnee, Peridot, muriat, lgrochow, JousterL, joy050530, ggaffield, dlsdl1210, Kfay, ninjew, minimalist, guygeva2311, orch, Sunnyway, cschein, Zeekza, TheAtom417, 1sigor, SBlanco, strangerxtheminute LynneC, wozzarelli, crazyman4865, Philippe_C, dragonturtle, smalljude, sambarko, dataminerstarr, SpookyGrowly, rightnet, cognaso, LeeMo127, sparkst3r, whitefieldcat, kinryuu, Phil98, eluchinat, sydnerdio, lobusparietalis, hewhoamareismyself randompersonjci, twotwos, wlindboe, r3, skruffylooter, papaiebiel, Mmalia12, LindseyAB, harkiesvf, SiliconOxide, baraitalo, sjapelson, ValdingerBella, juneyang2005, antarctica, Fourleaf37, Ayatee.

- Scythes helped correct reconstruction errors (number of corrected cubes): Nseraf (19141), susi (4116), galarun (1467), KrzysztofKruk (1453), nopasaran (1200), annkri (1136), scoobi (926), a5hm0r (863), aesanta1 (846), LotteryDiscountz (844), Kfay (816), rinda (715), Atani (639), ronin (428), dragonturtle (380), MaraTara (364), lilmoo (320), Baraka (286), crazyman4865 (258), Cliodhna (257), jinbean (251), toknow (240), jfine (232), michellewooten (229), nkem (219), ggreminder (186), kinryuu (178), tunerk7 (174), Caffeine (165), LynneC (164), m7md (159), b_hailey (146), skruffylooter (142), reb1618 (140), jax123 (139), alrianne (126), marhav (121), JousterL (101), addieye (76), baraitalo (70), Here_n_Now (66), aluorvats (65), DannyS (64), kronnn (60), kondor (54), hewhoamareismyself (54), eldendaf (52), Frosty (49), twister2 (49), jbinsc (48), Manni_Mammut (48), Andrearwen (44), bl4ckscor3 (38), Gruenewitwe (32), Oreliel (29), BladesRUS (29), Xnopyt (27), djajsl1234 (24), DannyScythe (23), nagilooh (19), tiikerikani (18), cognaso (18), tek50 (18), LindseyAB (15), Certh (15), Lunar_Dust (12), blackblues (11), SiliconOxide (11), randompersonjci (11), antoineb (10), twotwos (10), Fabiett (8), mam711 (7), r3 (7), mamateresa (7), dereks (7), sambob496 (6), chiflows (6), gardenpea (6), ian_danskin (6), davidjones1105 (5), aldof (5), st0ck53y (5), Emeraldstar (4), mcfly (3), iridium (3), blabbermouth (3), Oppen_heimer (3), twisterZ (2), hawaiisunfun (2), Talifan9 (2), giallocromo (2), minimalist (2), Braincrash (1), IcyCuber (1), klrtibov (1), CoolCream1232 (1), AzureJay (1), Noydor (1), wozzarelli (1), damocles357 (1), tickytocky (1), SpookyGrowly (1), rpeak1 (1), juneyang2005 (1), Snoopy4711 (1).
- KrzysztofKruk wrote add-on utilities and scripts, including code that improves the Scouts' Log, where Scouts and Scythes communicate with each other.

The 3697 EyeWirers who contributed to reconstructing starburst amacrine cells are (along with number of gameplay cubes):

Nseraf (87943), a5hm0r (66467), twister2 (56906), galarun (45087), susi (37477), jinbean (32570), iridium (27256), aldof (22898), jamiexq (20676), tek50 (20389), sdunn61781 (19066), rinda (18967), lobusparietalis (18119), aesanta1 (17840), MaraTara (16908), charleskoch (16874), mamateresa (16208), crazyman4865 (15639), benficus (15225), grizle (14968), tiikerikani (14112), sneakybaron (13377), Emeraldstar (13216), jbinsc (13041), damocles357 (11518), marhav (10308), sambob496 (10235), Begonie (10082), pkeoughan (9230), faunhaert (9068), LotteryDiscountz (8826), ronin (8305), ketta (8260), jcostantino (8086), DannyS (7828), reb1618 (7788), martinj (7732), 1sigor (7632), lynnschlos (7510), buco (7430), m7md (7177), michellewooten (7174), sappygoblin (6413), dereks (6408), sappy (6332), IcyCuber (6077), blackblues (5959), dataminerstarr (5642), b_hailey (5249), jesmith_nm (5093), jax123 (5054), keshlam2 (4994), lilmoo (4992), nspainter77 (4985), mardim2 (4696), Chainsaw_NL (4627), Cerebus (4547), dhill23 (4194), mmcdermott55 (4164), jyri (4150), eyegap (4147), sherryhamm (4101), couc (3981), melodiebenford (3910), carpwoman (3864), toocajun (3861), scoobi (3833), nopasaran (3739), peggyu (3727), chrisrolik (3633), lopezlv (3582), Atani (3495), Enma (3477), sjc2153 (3430), pistachio (3381), nkem (3374), mjyothik (3343), ccolbert (3334), ginny (3329), HariSeldon (3324), Laurcifer (3311), colleencat (3297), ericafulness (3281), clarrisani (3242), rogis1 (3190), wee (3116), toknow (2992), erinys (2883), psimka (2877), retepaskab (2842), davis3792 (2826), ic167 (2733), smalljude (2718), dejerpha (2642), tryffyd (2638), marika (2610), Andrearwen (2591), greglanders (2558), addieye (2506), rmissel (2479), JacquesG (2436), dbhunter (2416), Kekelktus (2392), nestorlo48 (2346), bdesmartis (2327), eboyle23 (2306), Cliodhna (2272), kinryuu (2257), jabsco (2220), mollica (2218), PhineasB (2218), bigbiff (2212), borrowedbluebox (2200), wlindboe (2186), uzenik (2185), shannonlevine (2144), beatrixbloxam (2141), ouiz (2096), twistedtime (2091), 17lion (2079), KrzysztofKruk (2070), giselachristine (2055), r3 (2049), dejavu1031 (2035), BlackCat13 (2000), npatnode (1997), tunerk7 (1995), wolfryder101 (1928), rutho13 (1919), dragonturtle (1911), anitram (1902), Lunar_Dust (1901), DELETED34474 (1800), judystarbuck (1792), eldendaf (1771), DELETED34638 (1710), mywire (1659), gruigah (1655), herty (1638), mhelm (1638), inockach (1631), lotus98 (1591), lili77 (1562), amyd508 (1558), Passiflora (1556), taboen (1544), graigrai (1540), cschein (1525), ssaba (1520), mileslane (1499), LynneC (1493), timoflowers (1492), lsw463 (1491), mariomar (1485), lcurtisadams (1467), fidel (1450), rillabee (1420), bemaline (1419), Kargoneth (1397), asd123 (1396), sarabc (1376), lisainsandy (1347), dougrike (1344), jsileo3 (1338), stitcher52 (1337), kloez (1333), twarning (1333), mbb480 (1331), Hannibal87 (1323), jh1109 (1321), aise (1311), mstanley (1311), newcity9141 (1310), vienna717 (1310), EllenRipley (1308), t2null (1305), sumo (1298), frabuleuse (1297), Quinlan. (1290), panosdalk (1288), bouncy70 (1284), ChelonianRiot (1265), lringelstetter (1251), tennifry77 (1246), kronnn (1245), pycospain (1232), jero (1223), coherent (1217), Pif (1198), argento92 (1178), kshearer (1166), Cat4248 (1154), ppotter613 (1153), soniac (1152), wojtekp (1151), Fritzchen (1146), furball13 (1123), whitefieldcat (1107), giallocromo (1101), hilmersbrain (1087), ren53nyc (1061), karenza (1056), besttry (1052), cadrake (1044), mmweiland (1029), alswns4097 (1027), dsart (1021), Lauri22 (1014), sjhy2050 (1013), schmelik (1007), vaeltaya (1005), pfenn (996), svincent (995), upcyclist (991), gartral (991), TheStatPow (990), batchen (987), rightnet (986), jfine (967), nagilooh (957), ohleyer (957), lyzzard (949), mrmathi (939), lukata (939), unburleyvable (926), aluorvats (922), Lgchinadragon (917), walty (917), echopapa (916), mam711 (910), ossomasticato (910), blabbermouth (904), ultrapeanut (896), PersonalGamer (896), vague_nomen (892), fofomazuzu (874), kirste27 (871), Varppi (866), Dandy (863), myrklv (859), jbarstad (854), baraitalo (852), Maejoh (852), otto (851), ArcanzaJenkins (849), FractalCuber (847), knaray1 (844), theo1977 (843), suburbanexile (834), yernagates (832), Philippe_C (831), brthelen90 (823), jslykhouse (811), Marta_M (808), bigredjeep007 (807), lexerific (806), es3ban (798), awakn (792), ssmtx6 (791), ssef0120 (790), laxcav (784), FluffySab (784), Mbrightjones (782), admiralfury (778), annmonty (777), bileduct (763), bmaco (760), auntdeen (753), eroush (751), Sujuperstar71 (748), tomoyo ichijouji (743), duckgu92 (743), bretfrd (737), amy (735), dryczko (735), jungmanp88 (727), Snoopy4711 (726), tlthornton (726), hank78 (726), lgrochow (716), marcus737 (716), garret85 (713), pishe (712), tschwach (712), Mastica (712), wozzarelli (704), mainbrain (703), co1ored (698), Manni_Mammut (690), jdoncarlos (687), Rosse (687), syllogic (686), catter (683), kab (682), sisterscience (681), sliet (679), Leopar (678), yagu44 (673), townshenge (670), jezoskoczek (669), muriat (667), skeeter_mcbee (663), qk2922qh (658), Unclewilley (655), pdksc (654), LaurieMarks (653), Mariahkitten

(652), cattrack (651), annkri (648), edremy (637), turtleshen (629), 000 (624), Civet (623), Iliyana (620), ducklette (612), hawaiisunfun (607), mkwak (605), mowens (603), adrian_alexis (600), gren25 (596), britishclimate (591), jonas.j.nordin (590), qkrw1 (590), elentari (590), jerryam (586), klimp (585), zuotian (581), yuval1 (579), annek (576), rekrab (571), jinsol6 (570), aebarnett1 (565), slycooper (560), engadin (560), AlWAl (558), deborahj (554), MissLiv (553), nasryn (550), mirandagavrin (550), generalnotes (548), gebe (546), teapackage (544), pandasecond (544), eluchinat (542), lazymuse (541), rigelan (537), bradtaylor (534), Sunnyway (534), tryeye (534), wjddls521 (531), DELETED14806 (527), chijunse (522), Christian_66 (522), christie (521), 0303sb (520), Gozgo (520), elliesaysmeow (518), sjw3310 (514), klaus13 (511), morphamagus (509), harkiesvf (508), redfish (507), maggielnb (505), jiberjabber666 (504), autograph (502), melissa0774 (499), supertiger (499), cksgh645 (499), sirpago (498), shamrockveg (494), Frosty (492), vickywu (491), dmlcks1228 (490), seu1140 (489), soulcollector (485), ablasky (485), Omoshne (483), honbioKGo (481), richardk (480), me (480), nosers (480), thrkajrek (477), summer7656 (475), annaorlando (473), hyungsuk4341 (468), kris3452 (468), Baraka (466), oc3711 (464), metamonkey (464), larobusto (463), kimjh0718 (459), janicemilliman (459), martyy_plp (456), VeggieNinja (455), kingboyd13 (455), ra3vdx (453), 01024984595 (451), rybaciagula (449), flyingrat42 (448), jw381097 (445), urs (444), Dodam (439), staso (439), cathy43dti (437), brandonberchtold (436), sheltiesmart (431), deadlyalgorithm (429), netic (427), pollysingerman (427), jogakxxx (425), cdaywalt66 (425), PWPeebs (424), Piepie (424), bunna (423), prohri (420), Fess (419), magic3033 (417), pduncan (415), yeehaw (415), qwenml (414), randomlogin (413), cooljohnny3 (413), dischordn8 (413), vezz12 (412), ngiaopao (408), awdzsx (407), kimjones1979 (401), corvidae (401), wurk (400), hyacinth04 (400), amyec (400), mcrisch (400), diizeiaj (399), seth (399), LindseyAB (399), dianasorg (398), mottiger (397), Cerialis (397), gedrod (396), kkim9932 (395), bookbird007 (395), jjw9706 (394), jjho9604 (394), Kfay (391), cls6679 (391), chiflows (391), willowbyrn (390), mh5052 (389), mdtheater21 (388), t3hnerevarine (387), alexmadsen1 (386), rudfbf4752 (383), kuhno1980 (382), minsak (381), karens (379), shadow_8472 (378), g1smo (377), ksup (376), noctivagus (370), mariemacdonald (370), hampton11235 (368), cptaaron (368), Altaire13 (368), gracemansfield (367), nashira (367), helge (366), Ceri (366), elise (366), flouf (366), iconforhire (365), Ditsch (363), neurosie (361), Hazelinka (360), LeeMo127 (359), Noeska (359), mind_less (358), cjg177 (358), kndahmen (356), Jem (354), momiga (354), LCU (353), Here_n_Now (349), mnbflute (349), jenwolfe8 (348), christsl (347), slated (346), redsoxwy (346), nj4876 (345), CathyK (345), rprentki (344), NoelleO (343), dkstkdgus328 (342), SpookyGrowly (342), dont_blink (341), dummypostfach (339), taceywhite (337), lookas (336), spoonbow (336), vilo_grey (332), ValdingerBella (331), ais (330), brianwilliamson (329), candychewning (329), caern (329), wolfspirit (328), jeoyoho (327), acida_2 (324), spriteling (323), wayner777 (320), Arthemis (320), carolmswiz (320), falconheart (320), readingite (319), pmh5954 (319), strangerxtheminute (319), duanestitt (319), hkc29 (318), alrianne (318), magret (317), lilbundlojoy (317), wrenoud (316), swoffler (315), miriamel (314), kwondsu (309), librarian (309), rkweir (307), JHGFD (306), berr_90 (304), titousensei (303), blissdish (303), canca (303), amblingpilgrim (302), awakaba (301), jmwc (301), ilona (301), logicality (299), dlwlstjr333 (299), haptic (297), blacat123 (297), lamebrain (295), marianne (295), YMT9999 (293), masterweaver (292), ccrowe (292), djajsl1234 (289), youn6501 (289), Nimrod (289), mcsnee (288), Pennywise (288), mck6 (288), siese (287), Fabiett (286),

Speeldinges (285), sungeun1214 (285), songbabe7 (284), machmuel (283), mimic751 (281), xCuriousCatx (281), skruffylooter (278), alexweasley (278), elch (276), brabender (272), gracewylu (271), robz90 (271), lvova (270), aaiken (270), kfailor (269), jmonderer20162 (269), ksleyk2 (269), frenchkl (269), Jlovin757 (269), lyndsey (267), kuzminski (266), kukumuzu (266), superdopamine (266), jardinpuzzles (265), heatherandjim (265), yoongsl (265), boomod (265), jte712 (264), Terramine (264), olive (263), ehdgml3391 (263), mrsmirrel (261), cengiz (260), paps (260), ItsNewToYou (260), plymouthdave (259), aliquot (259), tytoscope (258), maastolman (256), nicejune92 (256), rudska3443 (256), Ando (255), woaks223 (255), hewhoamareismyself (252), ghkdls99 (252), dellswanson (252), kenmierow (252), Vorlon (251), xodhks9205 (251), IsaiahC (250), Wendybird09 (250), dasjulchenm (250), Komorebi (249), stihial (246), 4x4jeepchick (245), Caffeine (244), Lilliscowleen (244), xpfjs1028 (243), kali13 (243), robertb (242), rkdanswns (242), ceh36 (242), secondlawlife (242), stevemj (241), 01085611037 (241), Richard_Henley (241), radeknek (241), mufintime (241), sht301 (239), hannele (236), wm05055 (236), sparkst3r (235), smos2022 (235), sommon4741 (234), deniceb87 (234), collurio229 (233), Jac (232), jordichanovas (232), mhutson (231), scopedriver (231), laurippt (230), sjfone (229), lieryan (228), lilyshield (228), honbiosra (228), Coconuts44 (227), RosDan (227), tunisiel (225), jenh (225), sbrooks_pilotmr (225), rankinc (222), angelas (222), mollyb (222), tobykenz1 (222), maxanti (222), Minokiller (219), clbriggs (219), senral (218), mirydian (217), hofstadter88 (217), wwwmkstr (217), ross_bales (215), meanae (215), bo2913 (213), guilifr (213), qorqoreh (213), Sekik (212), laccaria (211), tualilja (211), oroko (211), maxswiegand (211), notenbed (211), esbboston (209), efhw (209), raduban (208), lornetz (208), ramshorn (208), abotz (207), gadjo95 (207), ljk6431 (206), mberna00 (206), alessandracva (205), naspam (204), megaschara (203), authornd (202), ffffrank (202), flamingomarty (202), knzk (201), richarrr (201), vitadolce5 (200), ljy2715 (199), ishikakushin (199), Alya_N (196), profquad (195), catherinePouic (195), acidcats (195), bwstudent (194), omer (194), monarchspoison (194), AKhajah (193), chris71 (193), skl6284 (193), rkwhr3256 (192), susanmatthews (192), jorwat (191), Mereni (191), miriaml (191), mru (190), cmr17 (190), Magnoliahigh (190), 01059616163 (189), stlbluesfan46 (189), brinkhm (189), norm rhett (188), smilinghank (187), yev (186), honbioGAl (186), spaceoddity (186), mqrius (185), kamorge (185), solange (184), orko (184), withaar (184), lordzorax (184), theman2000 (183), mmichalka (182), randompersonjci (182), irene7micro (182), tjwns105 (181), TheAtom417 (181), lauli (181), Ania19 (181), hellisoued (180), wldndsla4 (180), jn291982 (179), Gwindor (179), ajemanuel (176), sambarko (176), LightPulse (176), whathecode (176), dms1044 (174), MaciekB (174), yeppi2002 (173), kastakan (173), annefallon (172), Xandrex (172), bjan (172), MikeGB (171), brianamywa (171), kseifnaraghi14 (171), newneuron (170), Sagaraghosa (170), steenezel (169), Ronnie_Soak (169), norbislv (169), alicia_11_11 (168), sfree4all (168), hiigaran (167), Lataru (167), codingTrickster (167), mahjong (167), sak3097 (166), jandi1203 (166), Saphira130 (166), laffers (165), hanes2002 (164), hi88hi (164), celiad (164), simonkaytamas (164), cestmarrant (164), qudgns0014 (164), mambosamba (163), gasavory (163), Freddo92 (163), CoolCream1232 (162), deliaknight (162), MissMilou (162), Bluerhyolite (161), seasharpe (161), hana03 (161), lkauth (160), pjoyce42 (160), kallahar (160), Kewne (159), whoheckhe (159), usagidark (158), zhfldk452 (158), wappentake (157), snail (157), BoredRaichu (157), EAG123 (157), tfrank (156), hharder (156), curious30004 (156), aimorai (155), shrajke (155), chase (155), jskiles (155), eee7016 (155), caphillipson (154), 5liwomir (154), deadxdying (153), davidheiserca (152), rosstamon (152), wallrue (152), pandabear (152), minimalist (152), darcpixie (152), woals3585 (151), ekiri22 (150), danube (150), nawaorl (149), StanleyScience (149), AlexPodmohylny (148), hanami (148), Artemis15 (148), louiscyphre (147), ybarthol (147), kwangyong810 (147), rlp1969 (147), Noydor (147), Jwb52z (147), dmacdougall (146), mjh1660 (146), slhszlls (146), physiocya (146), calico (145), alicestardust (145), jorge62 (145), serge (145), jsb54 (144), asnjhaley (144), Scot.a.murray (144), rae.heitkamp (144), elye (144), kjen (144), wendy529 (144), Sheepdog (144), bkahn (143), JoeNui (143), Crounus (143), RickGuar (142), petels (142), athensoh (142), ambermichael (142), auraseer (141), Manago (141), ns_neuron (141), rhazha (141), rvsobera (140), mdailey (140), dbarnkow (140), jwt1029 (140), Alfonso_P (140), mordodirosa (140), neuromancing (139), FMRTerrifc (139), maryal (139), lubaluft (139), devirumetmachina (138), JegElskerSnoeskyving (138), rutho (138), trembn2 (138), backupelk (137), won00789 (137), kamegani (137), hippo031 (137), bethleegy (137), peterahn (136), tmeekins (136), tracer911 (136), dilettard (135), maby (135), mlayten (135), dksehgus06 (135), thegreatgarbanzo (134), gdawg5130 (134), elev (134), pennymonger (134), chickadee76 (134), keross (134), Aaroncoyote (134), lookanelephant (133), siggenlh (133), ljk3053 (133), ehsansabri (132), markd21 (132), rafikh (131), MomHasAlz (131), SimoProvenzano (131), jjj78 (130), chadcoleman808 (129), coldfiregr (129), hibounoir (129), Cucumbersquirrel (129), rhfm123 (129), ggaffield (128), matias (128), seunghoaba (127), Azalais38 (127), mitja (127), kernfel (127), jpqj (127), rosapf (127), juneyang2005 (126), nemerle (126), irinayf (126), eragon1992 (126), wnsgur5443 (126), honbioNFu (125), mms502 (125), jojoful (125), 1129titon (125), HerDug (124), restcoser (124), emmask8s (124), dherkova (124), monet_open (124), knapper7 (124), cielreveur (123), Dr.Ron (123), pierceh2 (123), venerac (123), dwarshaw (123), lizbith20 (123), werecake (123), barpop (121), Mmalia12 (121), wksung (121), oliver007 (121), eselmer (121), geek2nurse (121), jkrzyz (121), Riley_Light (120), millenniumb (120), kasimclean (120), 65sprd (120), Geluemse (120), vlorbschnat (119), Rosiebud (119), muddart (119), turtlesee (119), leehy250 (119), yb0883 (119), nb6349 (119), nilinhim (118), zzahal (118), arachnae (118), aseo1106seo (118), sumizone (117), welo3 (117), jeffyap (117), salfordphil (117), Gryllz (116), cyjing (115), igor_chebanenko.72 (115), AbuMaia (114), david1908 (114), mapio (114), copantok (114), Waffle01 (114), Fhydra (114), Robadd (114), Danis_Bang (114), Wjebs (113), wormofsand (113), dswansonsb (113), webnajma (113), pagdzin (113), repate4444 (113), bobbygirl (112), dsprinzen (112), PrincessE (112), clarkphd (112), qwaszxopklnm (112), coryg308 (112), wjdiwh33 (111), EZM (111), luft (111), Flos (111), hklingon (111), jasperd (111), tommy1 (110), Disper (110), DodamTest (110), Neometron (110), skeptikitty (110), ikarus (110), lcallan (110), Cbay (110), gdnskye (109), joy050530 (109), kgweisman (109), valter (109), asdf3011 (109), jah (109), Ckarnish (108), 3rdibound (108), jaro87 (108), sagebrush (107), crewpet (107), good42n (107), cityghost (107), ecairns (107), beckeemo (107), JousterL (107), 8982679 (107), honbioSWo (106), OlgaUpyr_KS15 (106), medievalman30 (106), towever (105), THOREAU77 (105), chrysanthemum (105), kri5 (105), safetythird (105), andriejunas_lt (105), ggreminder (105), sheep1 (104), thajazzlady (104), summitpeak (104), kimgtea12 (104), chippervak (104), radon (104), junsung1250 (103), drbyte (103), Waltika (103), smoores (103), ppizzo (103), sidran (102), lolshanks (102), gadwicke (102), sandyilfs2 (102), penzor (102), michelle127 (102), orch (101), curiousimbroglio (101), parhaml (101), justagirl (101), ldm2020 (101),

01044645208 (101), tiatm (100), cyberdork33 (100), tndusl3000 (100), harmonicquirk (100), jujung27 (99), arzi1021 (99), magnuson (99), mcd8604 (99), roro78 (99), MaLo75 (99), crazyguy (98), heptolisk (98), tkatie217 (98), cjdeakin (98), papower1 (98), Sotigan (98), aprilwent (97), nietzsche (97), sophiagerje (97), nickid (97), skywalker1 (97), somisetty (97), DELETED75279 (96), fmerge (95), bikwai (95), JNakamura (95), yourion (95), tongo (95), 4A85E (95), whenpigsfly (94), Neoronenblitz (94), mjyo (94), viklar (94), pepramon (94), Shmease (93), McW (93), shackley (92), ghi06064 (92), arc330 (92), macdoes (91), derven (91), kwon3693 (91), dreadcat (91), Trumpeter95 (91), el_burnso (91), ibisenc (91), spunky11 (90), renegeneroso (90), guhi3 (90), Christophk116 (90), Bollo44 (90), MindBlown (90), mevlana (90), Exotje (89), dlwngh8720 (89), skreed (89), toscaxyz (89), Yark (89), Minazumi (89), shardana (88), burhop (88), IgorOhrimenko (88), drag2012 (88), kagej (88), lyndseyannette (87), care (87), annamatsen (87), invest09 (87), alanisfer (87), pleeze (87), Locutus (87), sandrad (86), kohlman (86), Hcom3 (86), marka (85), MamaDragon (85), msch.dk (85), chenzw (84), cake91 (84), omniczech (84), kevonni (84), potenti (84), golfluvr (84), kazarenko (83), xogus9698 (83), k2 (83), lindanicolette (83), serilleous (83), Ylandy (83), zing27 (82), orenico (82), DELETED273615 (82), uprudtjr123 (82), antoineb (82), emersoj (82), coconutO (81), shy960306 (81), Headofibis (80), wjhsiao (80), kuzvo240 (79), meitaka (79), sylverone (79), Bragsen (79), tamasi (79), edderiofer (79), thevogfather (79), czesiu (78), SarahAndIchigo (78), sassypants1126 (77), minisiebs (77), qkqjqkqj12 (77), erez (76), sukhyun99 (76), guygeva2311 (76), equus1556 (76), cocona (75), HollyinAshland (75), infinitebattery7 (75), mjviss (75), game0009 (75), moirasanna (75), gamid (75), koraynar (75), begin79 (74), VADER (74), Tatyana_Violet (74), cosmicdolphins (74), daisykho (74), colinra (73), grad2011 (73), egonzales (73), SBlanco (73), maxbgreenstein (73), rbtrry (73), Delco999 (73), semi_deluxe (73), asencio88 (73), Martinqua (72), bloekmedwe (72), Flocktime (72), mevi (72), Maja1986 (72), DELETED165909 (72), emusa (72), omen (71), TomMarcotte (71), andriejunas (71), tupperharley (71), trau (71), enavarrocu (71), greenlizard (71), janet (71), hsseung (71), seleniumsolace (70), cerebralcortex (70), turing8 (70), damian1307 (70), EricRoberts (70), revel (70), Gally (69), lemurs (69), thieum66 (69), vogon (69), michielrutjes (69), sethiroth66 (69), misterjstewart (69), story39348 (69), Vinara (69), xrnibor (68), GrandmereLouise (68), hitro89 (68), gentrymc (68), kjl3080 (68), DawningAventide (68), tarapratt (68), EichholzFan (68), glorioustan (68), mjeannez (68), noosey (68), emsabo (67), prowlerath (67), JeyTee (67), audreyreale (67), rosslh (66), darkchamp (66), physiosli (66), kobuki82 (66), Snibril (66), crystalhahn (66), anneli1221 (65), goodjelly (65), david.pavlicek (65), mferber (65), jeonging2 (65), poen (64), aostojic (64), vbperrotta (64), guarantee114 (64), Woong (64), pbailey (64), conrads (64), peter1986 (64), topspot (64), blowfish (64), chickleodeon (63), CobaltZeroni (63), mindcon (63), ancarius (63), superdave50 (63), cody.rice (62), Hotline (62), beagol (62), wlsgur7974 (62), Trefling (62), spes004 (61), trinity8036 (61), bschroe (61), adelso (61), wesdxc1998 (61), schnipper1 (61), tatscub (61), dragon7496 (60), 4df (60), moonsiri (60), Shapoval_Oleg_KS15 (60), anakonda1974 (60), educmale (60), ImAPotato (60), Ania89 (60), evanilssen (59), kjw2051182 (59), dlwjddns5 (59), JustHelping (59), tomtrussel (59), bbmt (59), kite86 (58), Kerdj (58), Tom_TBT (58), s0nia (58), limstory (58), malaclypse (58), Xnopyt (58), sciencegeek30 (58), xasapula (58), ohgoody (58), angie12321 (58), sorek.m (57), inapigs2eyes (57), entwirrer (57), georgina4 (57), aynmara (57), OrionWolf (57), sapphiresun (57), Emma_B (57), rhn (57), SusanCotton

(57), JacqR (57), lorach (57), kjm8738 (56), Ivet (56), prothus (56), kim20170 (56), srhjk (56), Wincey (56), sthomson06 (56), r12s (56), quetsch86 (56), boyan (56), _ (56), jfmoralesd (56), Zarkark (56), kiwichick (55), Aigh90 (55), tomiks (55), lizaperks (55), edoody28 (55), mattness (54), Calycaa (54), ashleyjames (54), Zane_628 (54), dkutas (54), deltapapa (54), read (54), Konafets (54), pascallev (53), heskey30 (53), DELETED (53), jzezel (53), ravissante (53), drand (53), Onhyro (53), waterbug123 (53), zyfi (53), yjyj1023 (53), SATLas (53), momadoc (52), KJG (52), hleisma (52), Phil98 (52), TheOnlyOne (52), phronimouse (52), hergot (52), billzhu (52), semaolvidao (52), etgst (52), hakeem82 (52), hillary625 (51), mneurocat9 (51), arrosa (51), ilu2forever (51), riddesh (51), Mayad070699 (51), shelby2190 (51), fabiodavilla (51), Teepfluecker (51), aspringer (51), empact (51), looker (50), thehoboclown (50), ary_star (50), djsketch (50), matollik (50), Antemmasia (50), elvetter01 (50), Zeekza (49), sjapelson (49), mpop (49), giovanna (49), szhangjerry (49), shinmei2006 (49), bebe1817 (49), rsp (49), happyyj1997 (49), hoursanov (49), ahanssenmn (48), cvanveen (48), carterch (48), Marval13 (48), jojefree (48), Springtrap_66 (48), chy1000 (48), physiorfl (48), lklockhart (48), cognaso (47), musca (47), jfpickard (47), mikearends (47), mjw063041 (47), Lochnivar (47), JohnnieDough (47), tobe_2098 (47), catywompus (47), gabba (47), nayoon0117 (47), ba1r0g (47), dlsdl1210 (46), shj00612 (46), lari_lh (46), lh2iwire (46), blancaratouli (46), sofpav (46), Zakhar_Krasnov (46), theresa123444 (46), kiksn (46), dewondolynia (46), nurtwalder (46), ARandomTool (45), elgato337 (45), echo (45), Randomfull9 (45), lunarhyane (45), Sallymose (45), brianna (45), louischarles90 (45), YCATS (45), twotwos (45), muh1117 (44), momos (44), n1k55 (44), swax (44), DigiAspie (44), kes2521 (44), demonom (44), bcnissley (44), scottwuzhear (44), ZonaCat (44), suneva (44), LEHoeger (43), brunered (43), hermes (43), Nyabi (43), hcmcminn (43), evel_chihuahua (43), jld592 (43), borgind1234 (43), uopo4859 (43), dracttosh (42), tomy515 (42), mcr103 (42), andante44 (42), null (42), Elcarim (42), Ma.K (42), gardenpea (42), avatazjoe (42), lemunet (42), Simbertto (42), sac_a_puce (42), yawnG (42), dowhatisay (42), jfgariepy (42), Albert_von_Herford (42), tyyppix (42), hepatica387 (41), deyanrashkov (41), Szelma (41), Sagheera (41), joshuajoakley (41), Swiper (41), kondor (41), Davhornfir (41), janjin (41), britl (41), Jamoni (41), eatmorepickles (41), Gritty_Reboot (40), gamez7 (40), renate (40), Taminnugget (40), jonesgirlferrell (40), kolalelf70 (40), hollyw210 (40), apmontagne (40), TPM_23 (40), snik (40), hl3mgr (40), anmrlp (39), zimmux (39), chocmoose (39), vsorana (39), cooooookies (39), reena (39), darajoe (39), lpsw0410 (39), compguy09 (39), mcglk (39), owenga (39), leraoesa (39), tajia789 (39), Queensalis (39), shpark2153 (39), gamerjars (39), McMasterpiece (39), DahliaBones (39), cnb24 (38), Rene.Torres (38), pupo1385 (38), 1nineteen (38), papaiebiel (38), tmdgus0577 (38), msully98 (38), MikeyG (38), jyjs1031 (38), ghanima (38), Falgoriend (37), Oguona (37), Dr.zippy (37), orionfyre (37), slasher (37), dansilber (37), baxisblue (37), skye2520 (37), kristachio (37), drawyks (37), sajeffe (37), funabout (37), lynn (37), freesong (37), ariverr (37), saeb101 (37), swan (36), paxon (36), sorgd (36), darkshadow44 (36), wiredanimal (36), artwinged (36), ekerstein (36), x9q9c (36), carleye (36), tets (36), jlambert (36), Byuda (36), kevers94 (36), fgyo (36), nisanick (36), jimmy_xcb (36), pinormous (36), mdmegill (35), DELETED168570 (35), ender2336 (35), bacon_bacon (35), Engine (35), jhulten (35), satoriend (35), trailofscales (35), pennb24601 (35), cedric (35), joac (35), claytonicus (35), GarrusVakarian (35), paethon (35), sciamanna (34), codecareerchange (34), Elenlinte (34), mataller (34), fenshome (34), df910105 (34), thepic (34), 1ong1ashes (34), Ererexiue (34), lemmert (34), jonnableel (34), astrocyte (34), learnedhappiness (34), tabby (34), Cmdr (33), Morphy (33), lordandmaster (33), Nomitratic (33), Maye (33), stingypanda (33), theneurax (33), erebus (33), zachboots (33), schjora (33), kwy9022 (33), curiousepic (33), Emirald (33), luxdiffusion (33), dmd76 (33), iskhakovt (33), davidmadsen (33), Anastasia1 (33), demorie1 (33), Peridot (33), mimimelo (33), ube2o (33), TomasMihalek (33), walkersister (32), GregO (32), gman (32), gemstate (32), PieThrower (32), lucas73 (32), haihami (32), platelmint (32), bluemchen (32), aruzan (32), Togo (31), aydin (31), 1206549 (31), Certh (31), mirandakarson (31), geologyrocks (31), anson1006 (31), sikan92 (31), FestinaLente (31), MagneticHammer (31), vincentadam87 (31), alou504 (30), jogarza (30), bugmenot (30), anueinberg (30), denisehigg (30), susansfb (30), physiojde (30), rksdlqkqh (30), sdenzler (30), ladykaro (30), Wow_such_neurons (30), gurdiac (29), amorvan (29), Tirla (29), smartina (29), leodavid (29), laurivan (29), deancolls (29), esashby (29), yanshan (29), gparker321 (29), cordi (29), nogoodkris (29), chris77f (29), akreid (28), jongmin2 (28), rickyspeak (28), acaceres (28), jimh (28), TheOne (28), dalbergaria (28), dltldud0 (28), jesiswinning (28), fracter (28), sueandarmani (28), timothynx (28), omorali (28), Plaidomatc (28), dv941294 (28), Chapmmac000 (28), tiwake (28), ch40z (28), hutch (28), tjd0413 (28), Mateutek (27), nekresh (27), ccampbell (27), maciek0899 (27), kevkingofthesea (27), lulugoesboom (27), AnaelG (27), mikecheney (27), tegal (27), amdou4 (27), guswn5828 (27), katja (27), carlosaag (27), Poovent (27), 4onen (27), charalampos86 (27), shibumi (26), mvdm1990 (26), it_ohs1018 (26), Richtefee (26), vehscle (26), lerler (26), polyenthusiast (26), arwenm (26), 1233803 (26), epistygne (26), mikkelsen (26), josiahmix (26), erhome00 (26), badams214 (26), tishow (26), physionti (26), aka1 (26), visualysis (26), Nemtsevp (26), shakey_man (26), baron (26), laxwolf (26), aco (26), jrj626 (26), ussmine (26), devotip (26), ristintin (26), zeneece (26), svedge (26), enoch07 (26), kubakoniec (25), Brasa (25), zombot (25), Dudelianer (25), Nibbler85 (25), DoraTrix (25), jmceballos (25), 22nd_Floor (25), 4nushaak (25), satsuma (25), jasonsmiegel (25), eyespy (25), Mewidy (25), Simulant (25), Bazhanova_o (25), karasutengu (25), alisacrisp (25), saskfire (25), rynmccarthy (25), charna (25), antto (25), etbebl (25), tommy (25), ChloemoShaw (25), coelacanths (25), prosequi (24), Sigmus (24), way_runner (24), devn (24), scidonk (24), computermage (24), stewhi12 (24), legilimensmaster (24), ctighe (24), jonacthon (24), superrogin (24), misterpriscilla (24), candygrrl (24), deamiter (24), dima (24), saxywolf (24), nsi04118 (24), macinenterprise (24), erdbeerknochen (24), HanJu (23), ree (23), jigumok (23), SuperNovaYay (23), minnu (23), icotton (23), Anthony_Lee_Rice (23), Talifan9 (23), netiad (23), 12261264 (23), Janner (23), emmet9796 (23), mishfish (23), seo646 (23), iuean28 (23), bradsquared (23), pursuit88 (23), nick_sadler (23), Deety42 (23), krvb (23), iam1105 (23), zzzz (22), claudiag_jou (22), whddms9269 (22), Anja66 (22), psh0944 (22), sherapchodron (22), gonadarian (22), DmitryMalkov (22), themany (22), claymossterrylee (22), artur.klauser (22), coch (22), synes (22), fairboxie (22), pjhgeneral (22), kkururu (22), maggie50 (22), limefox (22), njhinds95 (22), Alexej (22), Tomghc (22), Lizzie98075 (22), fiddlydoo (22), whaleback (22), Kaputnik (22), evin (22), alexetciboulette (22), LuaCat (22), quantum413 (22), sitravon (22), foxlock (21), owlgirl (21), brock (21), hunt (21), jasonpapacostas (21), Hundas (21), ewazlozlo (21), big_sasquatch (21), mchughes1 (21), HillObserver (21), sfarah1441 (21), annie3472 (21), idril.piesdeplata (21), kittiesgomeow (21), haleem0803 (21), hsureka3 (21), kpt_fox_trot (21), metalstorm (21), cassandra87 (21), preece (21), taryntelle (21), aetherean (21), calvinx (21), merlmoor (21), hjun9802 (20), eabrunelle (20), arlind (20), arcodetejo (20), gemmi (20), Napis (20), SendPizzaASAP (20), mihaib (20), mrjrgregory (20), urhuckleberry (20), hadonos (20), anmbia (20), RetinoidX (20), popeyez987 (20), gasper (20), jbridge (20), mysticflames25 (20), tawalu (20), alexgotsis (20), dani (20), envaal (20), Genoplex (20), harm1995 (20), aqbaq (20), Keiken (20), gusgraham (20), jalynx (20), mrnibbler (20), kcutshall0710 (20), Electricz0 (19), ... (19), jane.matsesha (19), jmpalevsky (19), corvus1 (19), saguti (19), Mateon1 (19), baldymyra (19), verderox (19), jessiej (19), kenardX (19), kockas (19), slslgld0 (19), kt1004972 (19), belle (19), box213 (19), Frankenshtine (19), xanthej (19), falhorn (19), gabhahn (19), Samaelle (19), seok4793 (19), silverfire (19), anyep (19), Fourleaf37 (19), odebowy (19), anglerfish (19), twyster1 (19), jbolland08 (19), Deathranger999 (19), newlunarrepublic (19), mm523 (19), fingerstyle (19), helmars (19), IRSmartKittyz (19), kellydh (19), davinjax (18), hunter3154 (18), TyggyToo (18), narvaezm (18), MoreInput (18), llanger (18), olti (18), zricker (18), thebarracuda (18), karijnk (18), vitsen (18), nashbery (18), Zaynee (18), miguel.jodra (18), devonjones (18), KilgoreTrout42 (18), doxikal (18), cronehaven (18), andane (18), ninjew (18), jontabaco (18), idatwell (18), GREERO (18), GoldenMela (18), ashartrand (18), gizan (18), acchilles (18), marius31 (18), d_asterisk (18), Falant (18), acrabb3 (18), graith (18), amandabluezzz (18), chocomofo33 (18), justdakota (18), edward2 (18), dr_alon (18), baileyspace (18), Palaing (18), bebopdaddy (18), hippydogmx (18), wayneis (18), bestanyol (18), fabag (18), CWCorrea (17), mosng (17), keyeri (17), andyaltom (17), toddsat (17), dngudxor (17), beef_simmons (17), doomed (17), sniperdgs (17), 14cucumbers (17), BananaChiu (17), nyxer (17), zekaiser (17), physiopch (17), dfbarnett (17), kraynor (17), 5.3decibels (17), Ring3r (17), jeantherapy (17), cenright (17), rprounds (17), lynnai (17), lucianom (17), mjlachman (17), rosalina (17), saracynical (17), physicspenguin4 (17), suimugen (17), albin (17), scrialex (17), physiobmu (17), azoarcus (17), smizr (16), beez (16), caitlina (16), mahankala (16), jujeelie (16), vetryk (16), Terrag (16), thinkertank (16), christiantaylor (16), korenm (16), cherip (16), tomtripp (16), brainitchy (16), gunnar (16), k_gulberg (16), jsm1310 (16), jaimeasbury (16), almightymatt19 (16), grimjar (16), sungwoo1230 (16), lorschur (16), thom001 (16), melannr13 (16), twibble (16), balkamm (16), greenbean (16), NeuroMS (16), rhdudqls98 (16), ashrai (16), Incubus_Mirage (16), thajokerwild (15), grahame (15), horakely (15), swyeon11 (15), danenglish (15), blueambit (15), def3ctive (15), kapastor (15), 101454 (15), gmilway (15), gahg (15), lunaluna (15), lauraw (15), JessicaS (15), patufet99 (15), PaulT (15), daximus (15), sprlm (15), moche (15), m0bb (15), royfongroy (15), michbell (15), claireoconnell (15), ceciliadolago (15), sajtos (15), opiboble (15), spagls (14), bentemjensen (14), Vivre (14), yoyojo721 (14), scndz (14), Zmei (14), ur23 (14), lambykins00 (14), optimusbryant (14), mariam (14), chaynlynk (14), tsandy (14), wjdduqn (14), stridesofcobalt (14), socialuprooting (14), anaikahas (14), sothe (14), ellietaylor (14), beammeup (14), rjzimmer (14), kisu (14), Gruenewitwe (14), ncutting (14), retribute (14), eriador (14), Psychlone (14), flask (14), apfenton (14), xaero (14), kartov (14), xyzavex (14), jurkov10 (14), kimchulho (14), brabant (14), HROXANNE (14), dcliffor (14), primadoepi (13), m4573r (13), pkly370 (13), mugzyrae (13), Chimichangas (13), tlbailey (13), maximilien (13), devad (13), stellarbrain (13), acmokeson (13), TinaS (13), nimbal (13), nfinitpls1 (13), wptaylor369 (13), Oppen_heimer (13), bigwell (13), alakaluf (13), FishKAA (13), chairflyer2002 (13), raunija (13), amadeus06 (13), pandragon

(13), xk66 (13), kittysu (13), nidor (13), sandpile (13), fuzic1 (13), jesup (13), hot6138 (13), hongcwoo (13), mzakk (13), jpeetz (13), Leenalee16 (13), perrypotter (13), smeden (13), kristin (13), Zagna (13), xyphink (13), tvidebaek (13), DELETED229995 (13), nathanielavery (13), bubbles82 (13), Nseraf1 (13), eyjaring (13), nec22 (13), triana (13), baronblack (13), emja (13), bytensky (13), midian (13), overk1ll (13), roukess (12), ringzhz (12), ash1111 (12), aj1139 (12), kkimtis (12), skaiskai (12), mrtaustin (12), feedfail (12), Griso (12), batooka (12), travsud22 (12), wanderratte (12), straus (12), LeonDaDude (12), jin01056 (12), Lethe (12), Shtern (12), peachbed625 (12), heatherupton (12), fortaaak (12), zyphyrus (12), erkab (12), dalcomlua (12), mayzor (12), kylef59 (12), tzarina8472 (12), gent (12), willschull (12), odl23 (12), jnpy (12), saspist (12), favrage (12), rookiemonster (12), conscience (12), gahh (12), zagra (12), allactaga (12), danielr1 (12), susan2012 (12), aggelito112 (12), korthalion (12), lovely (12), lovelybookdiva (12), outof (12), snake340 (12), abencomo14 (12), ninavelka (12), aaass28 (11), curious_one (11), xiaoju (11), lemoncustard (11), yanok35 (11), neko (11), BenSilverman (11), OliCol11 (11), jinseop (11), aanewalapal (11), astokes414 (11), VB (11), Tetrixia (11), jonyoull (11), kerhbu (11), DELETED49013 (11), karmidee (11), dunemi (11), jschetterer (11), davenjen99 (11), whatnow (11), anais (11), rypgall (11), bmirella (11), cescaspecs (11), skibumef (11), brandon_kirklen (11), presb (11), nicolasrenier (11), mordeckai (11), jidoux (11), salinas_smiles (11), sp4rt4n117 (11), crosax (11), Sam007 (11), Oberon4759 (11), graymatter_ (11), byledojutra (11), pcsteak23 (11), lquirola (11), Proteolisato (11), hermosa17 (11), frankielai (11), takashi (11), yew (11), dimension (11), hairyfotr (11), kfleitz (11), tomhughes (11), sid961 (11), dylan (11), dioneuty (11), lmhch56 (11), qj2130 (11), LegionMammal978 (11), TomServo (10), tommaso (10), Roboterbastler (10), vmd1972 (10), prang (10), shesfuntastic (10), ravenshadow (10), kkezir (10), staalk (10), silasyeem (10), tushon (10), ivanescu (10), bodrick (10), ph0x (10), yby6338 (10), jack241099 (10), randonneur (10), jgid17 (10), day145 (10), cogscimajor (10), true8wolf (10), jermainiac (10), bluey (10), verity (10), yottaflop (10), nb02531 (10), JSHolic (10), naro (10), pastelparasol (10), moustik01 (10), Galestro (10), joseerre (10), msblue (10), YellowVenom (10), l3erdnik (10), nursefg (10), spiral (10), juliedp (10), W1RE (10), livelife2 (10), zelnicki (10), headlong (10), Jpassionforknowledge (10), CLDP21 (10), dhobsd (10), kibouhi (10), loved (10), cialhcrb (10), mirabile (10), avrame (10), memery_uag (10), mstarr93 (10), mikebond007 (10), cuvier (10), ianislavus (10), nillaba (10), xy479why (10), azalea1011 (10), csomoanu87 (10), kyj981023 (10), mightyjongyo (10), priapil (10), Arenir (10), ladymac111 (10), cawiebalk (10), zkdlxh1532 (10), kisuklee (10), SkellyJelly (10), sudonym (10), meduolis (10), chrisadami (9), bipedal (9), edecota (9), cheesethief (9), marcorossi (9), jcsandlin (9), saldrian (9), Bibou (9), yoffset (9), smithweiss (9), sneider (9), Telizhenko (9), 136andon (9), KingSalah (9), denagorsheelan (9), Caprise (9), jeremyhansen (9), katerSonnbert (9), pablo01 (9), bellatrix (9), mlweiss10 (9), thickfreakness (9), birdycrazy (9), david113 (9), jvmonkeys (9), examinerPD (9), Spit (9), whatisthiseven (9), 31orcas (9), jlevy002 (9), jankins (9), zeljko (9), Nysha2 (9), mistic6314 (9), scooterholiday (9), ciaodejan (9), rebecca_m (9), Murk (9), kruton (9), mhs (9), bb_mke (9), ianh (9), Naduka (9), Boop (9), bernsh (9), doctor (9), eyewireorg (9), natxo (9), mesuno (9), sharoffv (9), morred (9), Miles349 (9), ajerick (9), Ariodante (9), RM (9), smokeditty (9), Caesium (8), meechl (8), Rumkin (8), DELETED3730 (8), Moltodor (8), sciencemsg (8), eydel94 (8), gwazdor (8), frost (8), sarahgraber (8), striver (8), kikkerlg (8), icewindhunter (8), m_fair1 (8), roditg (8), tots (8), OmegaDalek (8), manuel.freire (8), shuang (8), madidbel (8), metapharsical (8), GoBrazil11 (8), mctb32 (8), filipeyro (8), jkirvbro (8), she5los (8), elephantrx178 (8), Golaf (8), naitsirhc81 (8), ab020619 (8), jun105401 (8), jhdewitt (8), delsenno (8), cogolf (8), syoifczeri (8), arc6872 (8), montanubes (8), orange4boy (8), bruceware (8), Shaul (8), faithfind (8), asdfghjkl (8), eugenskt (8), tvalesky (8), trolske (8), thingy (8), cirbi (8), julienelson11 (8), bokutunoryu (8), lala360 (8), ldods (8), burlynate (8), mcwhitehouse (8), Creativity (8), mbconnor (8), daan1201 (8), tripp (8), mrfuturetense (8), sbgowin (8), paola2796 (8), thestaticstar (8), katling (8), ingeworm (8), ksslng (8), marisa0122 (8), keos16 (8), karajeank (8), MagnusDeLions (8), carlosryanez (8), Neonnq (8), critias (8), youn9023 (8), bcatt (8), brieser (8), Maggie1Mark (8), rufasu (8), feliciter (8), mbaldi72 (8), lun0807 (7), caraloopy (7), ozymerche (7), renato (7), rosieg (7), ren1234 (7), gitsum (7), cozyasian (7), shin (7), Chief220 (7), Nivox (7), Gman2004 (7), randomblahnomina (7), cyndic (7), chipbuddy (7), coffeyhaus (7), neo9 (7), LNorbert (7), mfmezz (7), twisocfan (7), minollo (7), girliegirl1991 (7), tobio (7), olim21 (7), silentnt (7), decompiled (7), jennya (7), leejiyeon7233 (7), lfouquette (7), limamoon (7), ulis (7), apg1985 (7), jekwon (7), killimanjaro (7), zephir (7), MadBlack (7), mistermole (7), dfsd23 (7), knorke (7), grizzlysquid (7), omgitsomeguy (7), Prism019 (7), danserig (7), vapu (7), darkangelx (7), reddu (7), andfal (7), opirnia (7), marialim (7), catgirl12466 (7), anaelle (7), doomyxx (7), richxs (7), anonymous563 (7), kingn8 (7), tigerpurple (7), JWC (7), sayaka (7), bush6984 (7), MadameZsaZSa (7), diegoveranc (7), apsydelon (7), brisance (7), yunyoung93 (7), Forager19 (7), whitefang3927 (7), tsien (7), zhyaxu (7), tool789789 (7), onemanrace (7), kshs67 (7), pknewbury (7), pewwpewpew (7), Janovic (7), aubrey (7), nadamucho (7), ezekielsharples (7), ic (7), igniskhan (7), eduardoleitao (7), vqtwtk37vg (7), paulw98 (7), mact (7), Stichflamme (7), karneithys (7), tamborilaire (7), clevermonkey (7), luuu (7), anexum (7), xiram (7), bltnole (7), veelckoo (7), Braineses (7), keithtastic (7), april487 (7), eloimartins (7), xyztu (7), Acgb (7), PRG3D (7), hckjk071012 (7), smerten2 (7), msfranceshan (7), jacobscaff (7), heinermann (6), simulacron3 (6), alymsin (6), impulse585 (6), isaac (6), hugolopes (6), Yeexzyz (6), spcorrea (6), broken (6), captainbrant (6), death (6), SlightlyMango (6), k80805 (6), cjricker (6), iona (6), rileyfranks (6), cdog (6), azeyetech (6), magiconline (6), blakboks (6), legu (6), nkaschwarzkopf (6), seldara (6), werebunny (6), llamedos4000 (6), juja (6), proline (6), misa_laree (6), Tata1993 (6), KaveyKaveMan (6), mkaz (6), jaxblazer (6), gabriemw (6), enki1729 (6), hexidecimal (6), Frenchman (6), eoin (6), serych (6), mythique (6), tau23 (6), ADE (6), Arlelka (6), patrick678086 (6), willettk (6), lustigbr (6), physioebe (6), adelka (6), miky617 (6), herr_jones (6), durandj (6), Wyimaginowana (6), dvixen (6), thenonsequitur (6), pewwer42 (6), alex791 (6), soccertigerman (6), MickyC13 (6), DELETED158295 (6), snogo (6), hamley (6), lpayne93 (6), entropy23 (6), beans (6), ejyuan (6), squand (6), joekintyhtt (6), Gud (6), palovoi (6), celestetsukino (6), bogdan1234 (6), Wulfh (6), pepito666 (6), alvin519 (6), kat646322 (6), shonali92 (6), grendelkhan (6), robillard2010 (6), kristin11 (6), Rattus (6), leilabaroudi (6), darcymarum (6), sallyp (6), ilariaaccarino (6), cube94 (6), dsimps (6), mas49 (6), uellp (6), Spem (6), jinxie2300 (6), romses (6), sunjess89 (6), mushufeeney (6), EdHolland (6), ayena (6), skalette (6), alexae6 (6), grumpy (6), hlapurocciel (6), gratreus (6), Maranca (6), jeremyvicencio (6), cornergoddess (6), thephantomracoon

(6), opop64 (6), meissnereffect (6), BlackPhoenix (5), caticat (5), dejan_dejan (5), joshuagenubath (5), kclairdelune (5), werebear (5), danis (5), kevin1kevin1k (5), donut (5), jfuller4 (5), daggsta (5), dyordan1 (5), bobby1818 (5), soffee (5), donis (5), ganbat (5), filipre (5), aemacleod (5), DELETED21003 (5), scottjones90 (5), rockcoinman (5), Voxelus (5), orbilliusIII (5), szemdrot (5), savagecelery (5), nawre (5), sydnerdio (5), asdavis1 (5), emiriku (5), pretera (5), tkdtb (5), sittles1 (5), nernio (5), kristasoltis (5), fsmizlord (5), goduck (5), montster27 (5), marygriff (5), Axy (5), jigsawmonster (5), gigiz (5), pedestrian (5), kane (5), FrankBlack (5), wolven_moonstone (5), gmulder (5), isagolden (5), kristin86 (5), mimom (5), sikkert (5), lubos.odraska (5), arobbins (5), ddil (5), mrswhich (5), Tassie (5), zenith (5), tarosic (5), jm3ndoza (5), wschalle (5), Timestark (5), phantomtigre (5), marty1885 (5), odobryaev (5), peterpezer (5), snowbear745 (5), zcaqd12 (5), moonillusion (5), jackrabbit (5), discordchild (5), gutmech (5), mschell (5), okto (5), mjz19910 (5), raphaelhamad (5), legrandchef (5), AlchemistCH (5), leia0207 (5), Gully (5), lutianyou (5), beholder (5), eyeguy2001 (5), ksmason71 (5), Luft1990 (5), dingles (5), diamond7 (5), ebourdon (5), daondaol (5), quincy85 (5), ferkal (5), ashleyporter (5), clewi (5), alexto300961 (5), pgitterman (5), 0din (5), lsprott (5), Lutinka (5), rharsh (5), frederic1 (5), shenguin (5), eoint (5), revcom (5), autumnzuz (5), timtimmytim (5), kcedie (5), whddnjs1234 (5), acamachodepass (5), CrashxD (5), steinman17 (5), morenci (5), ninanemila (4), digital_trauma (4), muellerseb (4), justhum (4), marchandise (4), sbricau1 (4), physiocon (4), redsanurse (4), gengf (4), kipmacsaigoren (4), wiseerni (4), joshwd (4), ragnaroknroll (4), danielsan90 (4), roieki (4), medeleine (4), shirtlessguy (4), Lensky (4), CountryHat (4), jungsicy (4), andrelaszlo (4), tkdnj88 (4), nkem_test (4), greydov (4), kkh20nice (4), cmbarnes (4), kwokandy (4), blak320 (4), atom12 (4), duskvill (4), alexys (4), deja (4), Tedux (4), zeptor (4), philcalhoun (4), phosphor (4), neiirap (4), epilady (4), haileyh (4), eg101 (4), Phosphorus.yt.and.zd (4), larsmars (4), venndiagrams (4), zode (4), zchan5 (4), desnoo (4), DCMP (4), jon_allan (4), magnificat (4), zespy (4), rnejako (4), hmiller (4), abassi896 (4), waterhyacinth (4), cgilbertmt (4), DELETED143081 (4), yumyum (4), Braincrash (4), and7sosa (4), pbrennan (4), Katniss333 (4), halo95945 (4), weatherdude (4), Radioactive7 (4), ranotamix (4), nivo (4), dubplate (4), gerrai (4), scotte (4), asulli7 (4), mzilber (4), admiralack (4), DELETED219575 (4), freezedancer (4), awoo (4), Kayarewhy (4), yowdeno (4), tjiannone (4), caller (4), phillip (4), hchinik (4), alyssa (4), jmaster (4), ktdame (4), osamc (4), Baltasar (4), imnikkib (4), rebeccajoan84 (4), danielmoncayo (4), heymoon (4), tkdtod123 (4), Jeff343 (4), tritonmars (4), lihwei (4), knw257 (4), miraborn (4), eudokia (4), superfrikkinman (4), T3rminallyCuri0us (4), kmgold (4), palmyra (4), huginn (4), blue_minded (4), galmix (4), abkfree1 (4), ssucoff (4), Volonden (4), cannibalnakki (4), zubbay (4), jacob5124 (4), mitchpf (4), reculeao (4), cherryberry (4), kate38 (4), bandg8t (4), nico13p (4), Rapha1 (4), davubu (4), benfold (4), arai (4), KDM17 (4), TheBigZocker (4), r (4), darinab (4), PillowStar (4), slimslowslider (4), neuro303 (4), Tallen01 (4), centein (4), Priit (4), hannahrcoles (4), anke (4), js_kim (4), mollyelliott (4), Mccara (4), 780tatia (4), nuno (4), marcialasvegas (3), kerry (3), aham (3), philrod94 (3), blacknovayzfr1 (3), blam417 (3), awcy (3), andyxd (3), andr345 (3), pierce (3), frankhold (3), josephthink (3), anars (3), whitetiger90 (3), michalj (3), dadon (3), Shadowvik (3), newplayer1234 (3), flank (3), mulan257 (3), juliancesar (3), blackbox (3), darkie (3), dfajar2 (3), azducky (3), lswitch (3), deathspire (3), cptorisya (3), Claudiagoestobagend (3), XebeX (3), laiyuekiu

(3), bankstoneditor (3), shmoo (3), a_pache (3), Malakiash (3), karach24 (3), njlae (3), lostnuttybar (3), toby1p (3), pop91 (3), kinglionheart (3), gos22 (3), kalacia (3), sbliven (3), parthcmodi (3), manentia (3), jmag (3), magdalenlaura (3), vipersrt3g (3), drlidija (3), hwhitesell (3), liverlover (3), sexypigeon (3), AlexFranco (3), schrambo (3), readper (3), kangarookim (3), siroderab (3), hailee (3), dlvt52 (3), sith (3), reubenbreeden (3), rossi_michelle13 (3), abrno (3), tracymoore (3), jumpkeh (3), caeboa (3), TerryVog (3), echo_2 (3), aaronmusick (3), mka121 (3), Colonel49 (3), viciouschicken (3), sean4046 (3), ztalat (3), deadhead14 (3), andrea_gonzalez (3), chanewell (3), barbara (3), nonagon (3), sousys (3), thorein (3), ozabluda (3), wallybeara (3), nastia_shkurenko (3), ruthal31 (3), jacboo (3), vivinew (3), yesluh (3), ladynatty (3), nefeli (3), avant (3), piesaregood2 (3), scrumpy (3), bjk (3), akamat (3), koshka_xvost (3), zeek50 (3), wiresepp (3), tmsodano (3), Michella92 (3), AcidAlchemy (3), calvis (3), imahar (3), sa_sky (3), susanl (3), sgtbird08 (3), tdmitch2 (3), ljw4741 (3), cormath (3), ooajayi (3), onlysonofjim (3), origamiraven (3), Stockbrot (3), guhwang (3), psiconico (3), chemistrob (3), richj (3), scohen211 (3), alfitoro (3), moe (3), smosmo (3), spatak1 (3), Bouchee (3), rumpole31 (3), paulolc (3), zzleedy (3), lllhv66 (3), rudwns168 (3), Hoodwinked (3), lgallowi (3), ciaran (3), jerouth (3), ozzjen (3), jwhyb (3), maiaperry (3), RezzeR (3), plurgh (3), lobalm (3), volvman (3), slender (3), steffenga90 (3), casevillan (3), martin.shue (3), evanshj (3), apostrophe_s (3), dixy0 (3), capibarbara (3), sooseok (3), platta (3), Bre7 (3), wonseochoi (3), aljen (3), pablopon (3), mon7468 (3), Frestil (3), krissykat13 (3), mans0930 (3), danielatamayoa (3), mDC (3), pliocene (3), mc_goa (3), fasarias (3), hyrsz (3), leper (3), aegoodin (3), minime123233 (3), brain_teaser (3), ki78765 (3), pierresolide (3), dambatenne (3), NekoYasaka (3), terato (3), herbertg (3), sinrotulos (3), naromi1 (3), lenny (3), pkwitch (3), rmadna (3), mk0208 (3), nevermind425 (3), alefisico (3), brad95411 (3), fasdap (3), Jhe (3), snake1989 (3), SaraMasarone (3), wombat (3), gmjowett (3), mawil (3), gollaber (3), erikito69 (3), pkwayn97 (3), LiuBang10 (2), mkantor (2), doctahjosie (2), marcot617 (2), rkril (2), gjtnwls0817 (2), pumpaton (2), gjguaman (2), ottavi (2), mcsnebber (2), mrcrois (2), lipovskyc1 (2), tikitija (2), mgirl726 (2), hahaid2003 (2), joy65 (2), rinza0100 (2), drh_w (2), napolitaj (2), mlsowers (2), magnos (2), minci_tw (2), lalekutluay (2), kylefish (2), lcsmith241 (2), kwill19 (2), kashmir1 (2), rlnhernandez (2), jpp1960 (2), minsuyang (2), omrial (2), ratloaf (2), ayoung27 (2), sjgedoc (2), H3A7P1 (2), Kingme13 (2), Joanna_Aldred (2), isethzenunim (2), familyfun (2), yikai (2), cozmictrigger (2), soonerborn (2), constipatedbunny (2), burzum (2), jarisman (2), RiftingFlotsam (2), Rocketman (2), Crostine (2), kiara (2), zhxst (2), agomar (2), mmonette (2), Dwarka (2), misgurnus (2), lagann (2), saltheart75 (2), eunoia (2), vicxy (2), cfindling (2), tara_a_stewart (2), lucas2001 (2), Alvin11 (2), crusaderv83 (2), shuvva (2), ecosan (2), mucmuc (2), gayle (2), musicalgal25 (2), Goldsternchen (2), ShotRock (2), Tlaloc_Creation (2), szahra (2), icepink (2), unclefritz (2), javafern (2), kd7937 (2), larwloszka (2), dreamevil (2), eversuhoshin (2), dloutan (2), sebast1an (2), Linceo (2), jtneu77 (2), ibul0000 (2), bdaumit (2), benji (2), rocketship08 (2), eliako35 (2), Tanaqalt (2), lostsamoan (2), mogueta (2), leandro (2), pixelthief (2), lemmiwinks (2), DELETED6150 (2), ritchiek (2), thecoolbro101 (2), ckho0777 (2), xnak62x (2), ajokewinks (2), sharkvajay (2), infinitekyle (2), FieryCuber (2), eyethethird (2), lpac (2), petiteagle (2), jorgito (2), gimpyy (2), yujin2125 (2), meecie (2), jackwhitescarver (2), seaside650 (2), ad2673 (2), dmg04158 (2), sirililja

(2), zkm (2), obscurusglacias (2), Prelok (2), emily14532 (2), jwblanchard (2), KandidKnight (2), dernub (2), zmajchek (2), tomas (2), snoodledumpling (2), Charlie1 (2), kimmyras (2), victorzale (2), imphobia (2), edoiks (2), eis271828 (2), bediko (2), Liuu (2), merus (2), texadam (2), GrimReaper_Stash (2), dobrochna (2), architect (2), muhoweb (2), nerfherder (2), ronno (2), muwave (2), quietcanadian (2), fluffykitty (2), bloodonourknees (2), legendch (2), mattatbat (2), interurban (2), brittanygs (2), joshwash (2), a11fred (2), SusanV8 (2), Auntecedent (2), Abfabjilly (2), darshan (2), rwalker89 (2), benwade (2), kkjhh6163 (2), amphy (2), wackelpudding (2), rodenj (2), TheGrinch (2), wesselinator (2), pacosafr (2), dokluch (2), rosbe880 (2), britgriggs (2), longhairedfreek (2), kchare2 (2), makayla.hawkins (2), luddeus (2), rahabg (2), euan466 (2), tracycooperjr (2), creays9203 (2), mwbtle (2), dominicbi (2), gvgriffin (2), tango1123 (2), katanabob (2), tlukko (2), bdr9 (2), cammmw (2), nymvaline (2), zlateskii (2), sineunju (2), gramps (2), sanjeether (2), skyqween (2), fibromyalgia (2), tutorialtest (2), stalwartspy (2), wolfoftheredrose (2), marianne1988 (2), jontti (2), potter76 (2), wisery (2), DannyScythe (2), gunedown (2), oliviadherrington (2), 246nat (2), eli.rue159 (2), tuffbin (2), tsioutsiouboom (2), gctompki0816 (2), dblboston (2), EngineerGamer (2), errrinnn (2), sisifolibre (2), gebauer (2), teknowlogysltns (2), jenk911 (2), Rainlover (1), sushi810 (1), geekboywonder (1), moabutah (1), pizzacato (1), rontogeny (1), wakebob (1), ashleinikkole (1), brinlong (1), derouinbill (1), Hokepoke (1), diggyb (1), Cdanhly (1), h.c. (1), richwalker (1), Gugfvfgv (1), RINERU (1), DELETED125798 (1), pattyg (1), qwackdragon (1), maleypjm (1), mlovell (1), kleviat (1), Rae_Glover (1), bosung (1), nikanna (1), frenesy (1), ekhower (1), gcole (1), redhooligan82 (1), drblood (1), datacute (1), albuddah (1), catfish (1), alrighttsss (1), Nocturn (1), sympheaceu (1), jed42 (1), danielsimpson (1), narmi29 (1), Bartt (1), cesiumfrog (1), vittorio99 (1), shesshe (1), phoebustam (1), roese0647 (1), majernickt2 (1), kokiito (1), hanm0 (1), nobu1015 (1), fripfrip (1), DUPPY (1), Cptnamurica (1), igodsman (1), lilygiraud (1), bersch (1), inaj39 (1), yihsuan1230 (1), jonathans59 (1), davidk (1), justintwayland (1), powertaurus (1), whomp (1), CameronFlegg (1), chbrchka (1), kandelinsky (1), GrimReaper (1), alicemargatroid (1), garam5716 (1), thekla (1), matrixcakes (1), aelscha (1), rachelly (1), hectorh (1), mkm26625 (1), captainscf (1), cherithegreat (1), jess7356 (1), Crabardaf (1), phus (1), turambar (1), viper (1), edwinshap (1), doesnofollow (1), collinz1 (1), cmahung (1), matkne14 (1), kathrynpeterson (1), zincdz (1), smaley (1), b1k2h3 (1), NodEngineer (1), Kamutkid (1), Nseraf2 (1), bowlerhatcabal (1), mfrost95 (1), kenziemac5392 (1), zombiphoenix (1), charly (1), spazzyorbit (1), neufronteer (1), mminaz (1), mpc75 (1), mitico (1), jyj5028 (1), nine13pm (1), IanBanks (1), SlopeDoctor (1), the_xrumor (1), lpierre (1), alysakow (1), e_cyril (1), tomberek (1), emane19 (1), miffed (1), tijh (1), vedrance (1), Madone6 (1), tuggernongy (1), Doro22 (1), songsmith (1), Ivanisevic (1), K_Raven (1), minion (1), madrone (1), thandley (1), gutgie_2010 (1), jsmidt (1), thorsten_mueller (1), jkononchikjr (1), cboanca (1), fowler.a (1), srballesteros (1), wickstrom33 (1), blaxalb (1), cfegpse (1), heking (1), Tesseract (1), brainmapper76 (1), Just_someone_curious (1), gianotts (1), emilythedork (1), blueraider (1), freshy (1), malcolmteschan (1), galarun2 (1), mourz (1), acheld (1), dsenette (1), nurdles (1), jonnoark (1), soldeen111 (1), jjecho (1), prigubert (1), samvdv (1), banas (1), kylemath (1), sladjana (1), rsonn (1), BrightLance900 (1), SOh (1), LunaSquee (1), JUNI (1), jjw1932 (1), tvanukas (1), pitty (1), enr987 (1), brennayard (1), bluegem (1), bestienne (1), sivis (1), jkafka (1), durzo (1), hjsender (1), PMuller (1), tmb86 (1), supeinjin83 (1), seharia (1), mmit (1), Landselur (1), lok (1), ctrhlik (1), warboss (1), rmarin (1), cybermycha (1), grehy (1), llin (1), tinkis (1), willgray (1), moekel (1), jujulu (1), mandaken (1), roastbeef (1), jmf028 (1), valacirca (1), atheismo (1), bos10blonde (1), weesie (1), gwalockey (1), TortoiseBucket (1), ybot (1), anginehb3 (1), nwarner (1), santenna (1), felixe (1), kartben (1), Olman (1), qwsazxc (1), bottem (1), cmillican (1), thetasanctuary (1), ethanruin (1), MooCakes (1), sophia.liu (1), laurenlabo (1), samantha7 (1), kmiller405 (1), iman (1), polis (1), mjvonstein (1), ravenlockhart (1), mtnutholme (1), bexchick (1), jabbott (1), Shadowflame (1), rykashin (1), catking42 (1), Samhyon (1), Tltltlt (1), bfarill (1), itamarhaber (1), conchacatalan (1), GeWilli (1), kixu (1), mimiheart (1), joe_h (1), lxhxdvm_1993 (1), soad90 (1), symbol (1), eoin_o_sullivan (1), jthiatt (1), Zuzeus (1), kevinwitak (1), tossefar (1), Jlarsm (1), wolfgray (1), Patarka (1), ashtest (1), Bormand (1), honbioAON (1), vfeigel (1), LD2 (1), littleblitz (1), roscoclibbins (1), tazchowmein (1), Uendel (1), sridharsm (1), KR_VaJil (1), Terpsichorean (1), Neuronix (1), cleonepetaotaku (1), MisaLaree (1), greysky (1), theshortearedowl (1), neuroptics (1), mtvlukas (1), kkrr (1), orcmando (1), jkmitche (1), myth (1), rivka (1), fdgdfg (1), prm1016 (1), Juddric (1), Jimbo12 (1), andrejkelemen (1), mia_tsang (1), treyher (1), duncan50 (1), sabs (1), kacyk (1), Pat0alex (1), mauriciod73 (1), Daria_S (1), jvaughn (1), jnorman98 (1), atolby (1), sarkata (1), nszeihen (1), jtberz (1), btel (1), built (1), spiclette31 (1), starlightjumper (1), fernand0andrade (1), ddung0719 (1), pieceofjelly (1), JellyWrath (1), slysamdogs2 (1), dfrank_a (1), EndWorld (1), pandemonium (1), viscerator (1), Mio30 (1), weaselatnature (1), zaidimasakis (1), istarin (1), NONOSORRY (1), kintrbr (1), fletcheh (1), ltdrum (1), neevin (1), saksoft2 (1), Kilrathi (1), georgezgf (1), yeti_boy (1), dysp (1), lynni_aeon (1), czager (1), Del9fina (1), minase (1), skunk12 (1), huhe (1), edwin (1), dgdg (1), derslyr (1), ihaveregrets (1), evagrace (1), dagillo (1), CTRjfl (1), sybilmcshane (1), ovan (1), testtesttesttesttest (1), 3rnieXL (1), mullintj (1), hanaa (1), reo468 (1), ka9dgx (1), MandelDoom (1), tycdum (1), alta080 (1), picoos (1), aqwapengwan (1), johnmaxwell22 (1), rosie95 (1), doping (1), kn8aj (1), fizix (1), artic (1), jpouchly (1), 10562034511j (1), winta641 (1), BladesRUS (1), mattfolkerts (1), moskaluk (1), Moonwatcher (1), sini (1), molord (1), andyinchicago (1), lemurrr (1), brockwayl (1), migueldc (1), netsash (1), cjy161 (1), s210798 (1), shadowcross (1), tylas (1), mbkjer (1), lycorain (1), inco (1), freethesouls (1), Irthene (1), Torros (1), ggeu (1), bevalorous (1), helen5 (1), izziey (1), bytor (1), dalgaier (1), gpbabineau (1), Mister.MIke.reed (1), bliss25 (1), ricrdohdz (1), trabyl (1), jacksplay (1), lupusfavst (1), caitlin219 (1), johntomronsonbo (1), hohojojo (1), jimlaurent (1), liam_b82 (1), balkamm3 (1), jojjojojje (1), valarywithawhy (1), vidawells (1), slowjam1993 (1), tintintin (1), hesperia17 (1), jkeegstra (1), linkceo (1), sharksting (1), angrywolf (1), twister (1), dpath1015 (1), simkus (1), jaume (1), smithsam163 (1), flan54 (1), 130n (1), ferusvitae (1), brigl (1), kingzan (1), ayush (1), CharlieLeeLee (1), waterlord (1), ignolan (1), KlausiMausi (1), midashand (1), Vanderdoug (1), Oreliel (1), nadavy (1), highclass (1), hongjoohong (1), brainythebrain (1), 01083706775 (1), pheonixblade9 (1), diamond_dust (1), meeki (1), nomnomcandytime (1), lunasfav (1), gabgolly (1), anonymous929 (1), sloace29 (1), kyriakk (1), mtammeraja (1), anellium (1), xiphopagus (1), katvan (1).
